# Using cGMP analogues to modulate photoreceptor light sensitivity: Perspectives for the treatment of retinal degeneration

**DOI:** 10.1101/2022.02.07.478618

**Authors:** Sophie Wucherpfennig, Wadood Haq, Valerie Popp, Sandeep Kesh, Soumyaparna Das, Christian Melle, Andreas Rentsch, Frank Schwede, François Paquet-Durand, Vasilica Nache

## Abstract

Cyclic nucleotide-gated (CNG) channels play an essential role within the phototransduction cascade in vertebrates. Although rod and cone light responses are mediated through similar pathways, each photoreceptor type relies on a specific CNG-channel isoform. In many forms of retinal degeneration, increased cGMP levels initiate a pathophysiological rollercoaster, which starts with an enhanced CNG-channel activation, often in rod photoreceptors. This causes cell death of both rods and cones, and eventually leads to complete blindness. Unfortunately, the targeting of the desired channel isoform still constitutes the bottleneck for many therapeutic schemes.

Here, we present a novel strategy, based on cGMP analogues with opposite types of action, which allows for the selective modulation of either rod or cone photoreceptors. A combined treatment with a weak rod-selective CNG-channel inhibitor (Rp-8-Br-PET-cGMPS) and a cone-selective CNG-channel activator (8-pCPT-cGMP) preserved the normal CNG-channel function at physiological and pathological cGMP levels. The effectiveness of this approach was tested and confirmed in explanted mouse retina. Under physiological conditions, the inhibitor silenced the rods selectively and decreased the dependency of cone responses on light intensity. Remarkably, the activator, when applied together with the inhibitor, reinstated only the light responsiveness of cones. Yet, when applied alone, the activator dampened rod responses more strongly than those of cones.

Hence, combinations of cGMP analogues with desired properties may elegantly address the isoform-specificity problem in future pharmacological therapies. Beyond therapies for retinal degeneration diseases, treatments based on this strategy may allow modulation of visual performance in certain light environments or disease conditions.

**One Sentence Summary:** A combination of cGMP analogues with opposite types of action can selectively modulate either rod or cone photoreceptor function.

## INTRODUCTION

The phototransduction cascade takes place in photoreceptor outer segments and undergoes similar steps in rod and cone photoreceptors. To successfully fulfill their function in dim-light and bright-light conditions, respectively, rods and cones possess a complex photoreceptor type-specific cluster of proteins that is responsible for their distinct functional properties (*e.g*. light sensitivity, reaction speed, light-dependent adaptation capability, *etc*.) (*1, 2*). A key player in the phototransduction cascade is the cyclic nucleotide-gated (CNG) channel, which triggers the so-called “dark current” in photoreceptors. In the absence of light, CNG channels are kept open by high cGMP levels, which are estimated to be ~3-5 μM in rod photoreceptors (*3*). Upon illumination, rhodopsin becomes active and sets in motion a cascade of events leading to a decrease in the cGMP concentration. As a result, CNG channels close, causing membrane hyperpolarization and an abrupt decrease in synaptic transmitter release (*4, 5*).

Rod and cone CNG channels differ not only in their subunit composition but also in their functional characteristics (*5*). These channels consist of CNGA- and CNGB-type subunits (*6–8*). The rod CNGA1 and cone CNGA3 subunits confer the main channel properties and can form functional homotetrameric channels in heterologous expression systems. The modulatory subunits, rod CNGB1a and cone CNGB3, are not able to form functional homotetrameric channels, but play a decisive role for membrane trafficking and channel modulation by Ca^2+^/Calmodulin (*9–12*). Moreover, differences between the CNG-channel isoforms were reported regarding their ion permeation, ligand sensitivity, gating kinetics of activation and deactivation (*5*).

Inherited retinal degenerative diseases have a major impact on patients’ daily lives, compromising vision, arguably the most important of human senses. This is typically caused by the degeneration of either rod photoreceptors, as in the case of *retinitis pigmentosa* (RP) (*13, 14*), or cone photoreceptors, as in the case of *achromatopsia* (*15*). In RP the primary loss of rod photoreceptors is followed by a secondary degeneration of cones, leading to severe visual impairment and ultimately blindness, with no efficient cure so far (*16*). RP is genetically very heterogeneous, making the development of a common therapy for all forms of the disease very challenging (*17*). A fact that is well agreed upon is an increase of cGMP levels in affected photoreceptors (*16, 18–20*), yet the pathway by which the cGMP-homeostasis imbalance leads to photoreceptor death is still unclear. The most important cellular targets for elevated cGMP are the CNG channels and cGMP-dependent protein kinase (PKG) (*21*). In a study on the *rd1*-mouse model for RP lacking the phosphodiesterase PDE6 β-subunit, degeneration of rods was attributed to the massive opening of CNG channels and the subsequent Ca^2+^-increase (*22*). Moreover, when membrane incorporation of rod CNG channels is hindered by deletion of the CNGB1a subunit, retinal degeneration in the *rd1* mouse was significantly delayed (*23*). Similarly, exacerbated activation of PKG has been shown to cause cell death in several cell types and RP-animal models (*24–26*).

Based on these findings, cGMP-signaling became a promising target for pharmaceutical approaches (*27*). Several on-going strategies aim to delay rod death so as to prevent the secondary loss of cone photoreceptors (*28, 29*). Recently, it was shown that the photoreceptor degeneration can successfully be delayed by means of inhibitory Rp-configurated phosphorothioate-modified cGMP analogues. In particular, Rp-8-Br-PET-cGMPS (CN03) robustly protected rod photoreceptors and preserved cone photoreceptor function in three different RP-mouse models (*27*). The effectivness of this treatment was attributed to the simulteous inhibition of both high-cGMP targets: the “Rp”-modification to cGMP typically inhibits PKG (*30*), while the β-phenyl-1,N^2^-etheno-modification (PET) inhibits CNG channels (*31*). However, the direct effect of this and other cGMP analogues on rod and cone CNG-channel isoforms and whether they might be channel-isoform selective is unknown. While indiscriminate inhibition of CNG channels might prevent rod degeneration, it would also result in loss of cone-mediated vision in patients. Therefore, a pharmaceutical treatment should target only the desired photoreceptor and CNG-channel isoform, without hindering the function of the respective other photoreceptor type. Unfortunatelly, to date there are no known selective cGMP- or cAMP-analogues that can differentiate between various CNG-channel isoforms.

The aim of this study therefore was to identify cGMP analogues that could selectively target the CNG-channel isoform of interest, such that in RP only the pathologically increased activity of rod channels would be inhibited. While we were unable to identify a seletive rod-channel inhibitor, we found a selective cone-channel activator. By combining a general CNG-channel inhibitor with a cone-channel activator, we were able to inhibit rod channels and preserve cone-channel function, under pathological, RP-like cGMP concentrations. The effectiveness and extraordinary flexibility of this approach was tested first on CNG channels expressed in *Xenopus laevis* oocytes and further confirmed by micro electroretinography (μERG) recording on wild-type mouse retina.

## RESULTS

### Rp-8-Br-PET-cGMPS reduces rod and cone CNG-channel activity

cGMP analogues have previously been shown to be efficient modulators of PKG and CNG channels and/or to have neuroprotective effects in RP-animal models (*27, 30–32*). Here, we tested ten cGMP analogues, including several novel compounds, for their effects on rod and cone CNG channels (Fig. 1). To assess the selectivity potential of these compounds, the respective protein isoforms were heterologously expressed in *Xenopus laevis* oocytes. The oocytes co-injected with RNA coding for CNGA- and CNGB-subunit types expressed heterotetrameric channels composed of CNGA1 and CNGB1a subunits in case of rods and of CNGA3 and CNGB3 subunits in case of cones (Fig. S1). These channels exhibited L-cis-diltiazem blocking and cAMP-activation properties similar to native CNG channels (*5, 7, 33–35*).

**Fig. 1.**
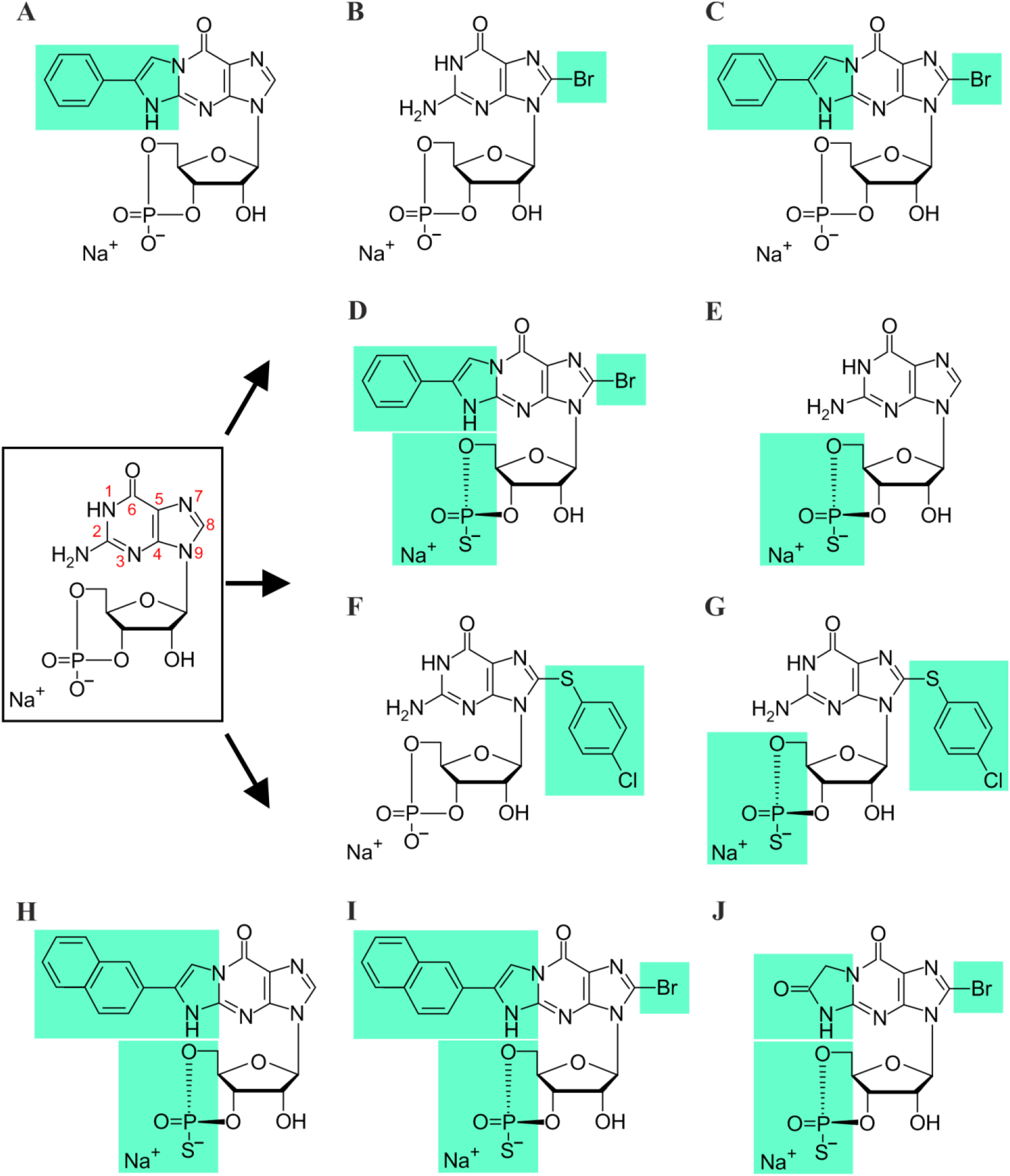
Chemical structures of cGMP and its analogues: **A**) PET-cGMP, **B**) 8-Br-cGMP, **C**) 8-Br-PET-cGMP, **D**) Rp-8-Br-PET-cGMPS, **E**) RP-cGMPS, **F**) 8-pCPT-cGMP, **G**) Rp-8-pCPT-cGMPS, **H**) Rp-(2-N)ET-cGMPS, **I**) Rp-8-Br-(2-N)ET-cGMPS, and **J**) Rp-β-1,N^2^-Ac-8-Br-cGMPS. Structure of cGMP is shown in the black box. Green background indicates modifications to the cGMP molecule.

For a better understanding of the effect of the Rp-configurated phosphorothioate-modified cGMP analogue Rp-8-Br-PET-cGMPS (*27*) on retinal CNG channels, we determined first the influence of each of its modifications. cGMP with bromine at position 8 of the guanine-ring system (8-Br-cGMP) had similar efficacy but a higher potency than cGMP, when activating the retinal CNG channels (Fig. 2A-D). In contrast, cGMP with the spanning β-phenyletheno-modification (PET) at positions N1 and C2 (PET-cGMP) behaved as a partial agonist (Fig. 2A,B). Surprisingly, the presence of both modifications in 8-Br-PET-cGMP reduced the efficacy to activate the channels even more (*e.g*. for rod channel to less than 1% of maximal activation). A similar effect was observed also for the Rp-8-Br-PET-cGMPS.

**Fig. 2.**
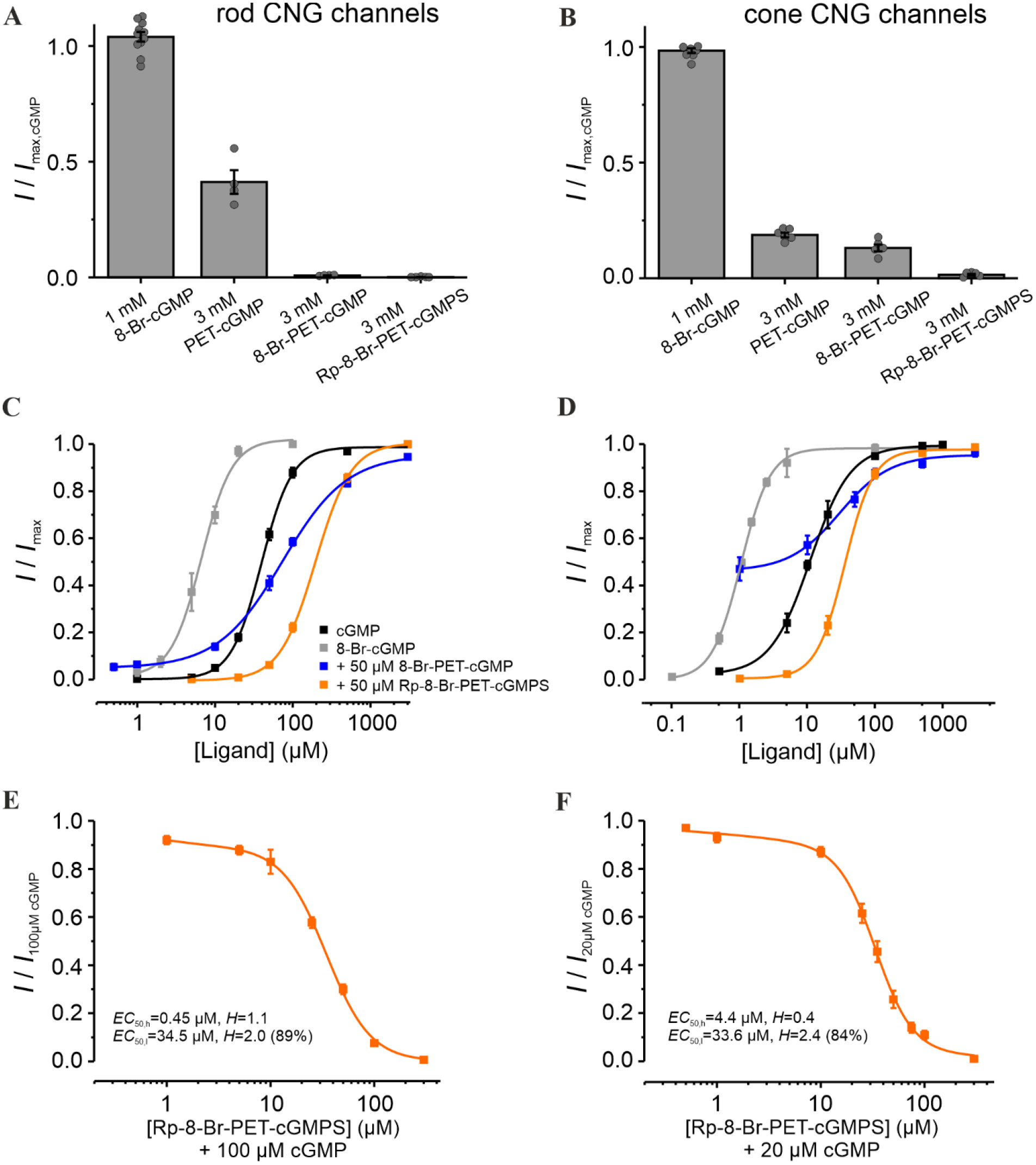
Effect of Rp-8-Br-PET-cGMPS and related cGMP analogues on retinal CNG channels. **A,B)** Efficacy of 8-Br-cGMP, PET-cGMP, 8-Br-PET-cGMP and Rp-8-Br-PET-cGMPS when activating rod and cone CNG channels. The currents measured at +100 mV, in the presence of either 1 mM 8-Br-cGMP or 3 mM of the other cGMP analogues, were related to the maximal current induced by saturating cGMP (3 mM for rod and 1 mM for cone CNG channels; n = 4-11). The grey symbols represent individual measurements. **C,D**) Concentration-activation relationships for rod and cone CNG channels in the presence of cGMP (black), 8-Br-cGMP (grey), cGMP + 8-Br-PET-cGMP (50 μM, blue) and cGMP + Rp-8-Br-PET-cGMPS (50 μM, orange). The data points representing means of several experiments were fitted with Eq. (1). **E,F**) Rp-8-Br-PET-cGMPS-inhibitory effect on CNG channels in the presence of 100 μM and 20 μM cGMP for rod and cone CNG channels, respectively. The experimental data points representing means of several measurements were fitted with Eq. (2) (n = 8-16 for rod; n = 5-12 for cone; for *EC*_50_, *H* and n see Table S1).

To simulate the strong variations of cGMP levels in photoreceptors, especially under RP-like conditions, we measured the effect of the respective cGMP analogues at different cGMP concentrations (Fig. 2C,D). Rp-8-Br-PET-cGMPS (50μM) led to a decrease in the apparent affinity of CNG channels: ~4.9 fold and ~3.2 fold for rods and cones, respectively. Thus, under these conditions, no isoform selectivity was observed. Interestingly, 8-Br-PET-cGMP had a dual effect: under physiological cGMP it produced a potentiation of retinal CNG-channel activity, whereas at higher cGMP it caused minor inhibition.

To determine the maximal inhibitory potential of Rp-8-Br-PET-cGMPS and to compare its effect on cone and rod CNG channels, additional measurements in the presence of constant cGMP triggering ~80% channel activation (*i.e*. 100 μM cGMP for rod and 20 μM for cone channel) were recorded (Fig. 2E,F). Below 10 μM Rp-8-Br-PET-cGMPS, the concentration of half-maximal inhibition (*EC*_50,h_) for rod channels was ~10 times smaller than that for cones (0.45 μM *vs*. 4.4 μM). Above 10 μM, the concentration of half-maximal inhibition (*EC*_50,l_) was similar for both CNG channel types. These results suggest a weak concentration-dependent selectivity of Rp-8-Br-PET-cGMPS for rod over cone CNG channels at low inhibitor concentrations.

### 8-pCPT-cGMP shows a concentration-dependent selectivity for cone over rod CNG channels

We next studied the effect of Rp-8-pCPT-cGMPS and of its relative, 8-pCPT-cGMP, *i.e*. cGMP analogues with a “4-chlorophenylthio”-modification (“pCPT”) at position 8 of the guanine ring system. Compared to cGMP, 8-pCPT-cGMP had similar efficacy in opening retinal CNG channels, but a much higher potency (~ 63 times for rod and ~138 times for cone channel; Fig. 3 and Table S1). Noteworthy is the robust difference in the apparent affinity of 8-pCPT-cGMP: 0.08 μM for cone *vs*. 0.63 μM for rod CNG channels. Hence, this compound may enable a selective activation of cone CNG channels. Addition of the “Rp“-modification to 8-pCPT-cGMP, in Rp-8-pCPT-cGMPS, drastically reduced the efficacy of this compound to activate both CNG-channel isoforms (Fig. 3A,B). These observations differ from those of a previous study where Rp-8-pCPT-cGMPS was estimated to trigger almost ~93% of maximal CNG-channel activation in rods (*36*). When the cGMP-activated channels were exposed to 50 μM Rp-8-pCPT-cGMPS, their activity at physiological cGMP increased and at higher cGMP levels decreased for both channel isoforms. Due to the strong increase in cone-channel activity at physiological cGMP, Rp-8-pCPT-cGMPS was not considered for further studies.

**Fig. 3.**
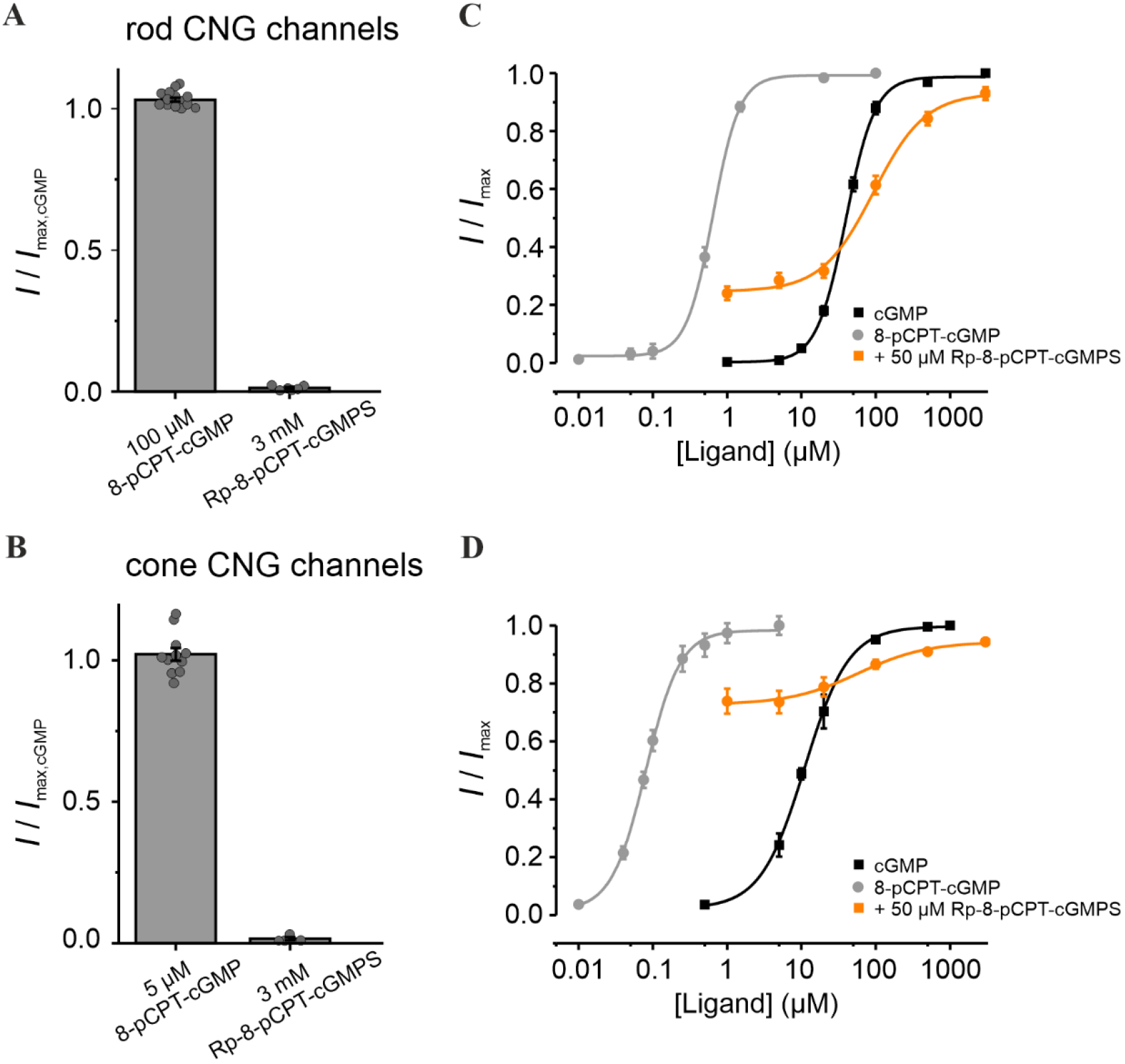
Effect of Rp-8-pCPT-cGMPS and 8-pCPT-cGMP on retinal CNG channels. **A,B)** Efficacy of Rp-8-Br-pCPT-cGMPS and 8-pCPT-cGMP when activating rod and cone CNG channels. The CNG-channel currents triggered by the respective cGMP analogues were related to the maximal cGMP-induced current measured at +100 mV (3 mM and 1 mM cGMP for rod and cone channels, respectively; n = 4-16). The grey symbols represent individual measurements. **C,D)** Concentration-activation relationships for rod and cone CNG channels obtained in the presence of cGMP (black), 8-pCPT-cGMP (grey) and cGMP + Rp-8-pCPT-cGMPS (50 μM, orange). The experimental data points, representing means of several measurements were fitted with Eq. (1) (for *EC*_50_, *H* and n see Table S1).

### Rp-modified cGMP analogues are not selective for rod or cone CNG channels

We next tested four additional Rp-modified cGMP analogues: Rp-cGMPS, Rp-8-Br-(2-N)ET-cGMPS, Rp-(2-N)ET-cGMPS, and Rp-ß-1,N^2^-Ac-Br-cGMPS. These compounds had a lower efficacy than cGMP in opening the retinal CNG channels (Fig. S2A,B). Moreover, except for Rp-cGMPS and Rp-ß-1,N^2^-Ac-Br-cGMPS, the cGMP analogues shifted the cGMP-induced concentration-activation relationship of rod and cone CNG channels toward higher cGMP levels (Fig. S2C,D). Although in the context of RP-type diseases this effect is desirable for rod channels, the reduction of cone CNG-channel activity at physiological cGMP levels would be detrimental to vision. Rp-cGMPS and Rp-ß-1,N^2^-Ac-Br-cGMPS did not influence the activity of cone channels but, unfortunately, they were also not isoform selective. When comparing the *EC*_50_-values obtained for all cGMP analogues, we concluded that, none of the compounds had selectivity potential for a specific CNG-channel isoform (Fig. S2E,F). Noteworthy is that Rp-8-Br-PET-cGMPS, the cGMP analogue which showed a protective effect in RP-animal models (*27*), caused the smallest decrease of the channels apparent affinity in cones, while the *EC*_50_-value of the rod CNG channel was considerably increased (up to ~5 times; dotted box in Fig. S2E,F).

### Combination of 8-pCPT-cGMP and Rp-8-Br-PET-cGMPS preserves rod and cone CNG-channel function under RP-like conditions

In an attempt to inhibit rod CNG channels, while preserving cone CNG-channel activity, we combined the weak rod-selective inhibitor Rp-8-Br-PET-cGMPS (the “inhibitor”) with the strong cone-selective, potent agonist 8-pCPT-cGMP (the “activator”). We studied the CNG-channel activity in the presence of cGMP, inhibitor, and activator, and compared it with the one triggered by either cGMP alone or cGMP and inhibitor only (Fig. 4). Several activator concentrations around 0.1 μM (*i.e*. 0.2 μM and 0.05 μM) were tested, because this concentration triggered ~90% cone CNG-channel activation, with almost no effect on rod channels (Fig. 3C,D). The selectivity potential of this treatment was estimated at cGMP levels likely to occur in early RP: for cones at “physiological” cGMP (~5 μM) and for rods at pathologically high cGMP levels (~100 μM).

**Fig. 4.**
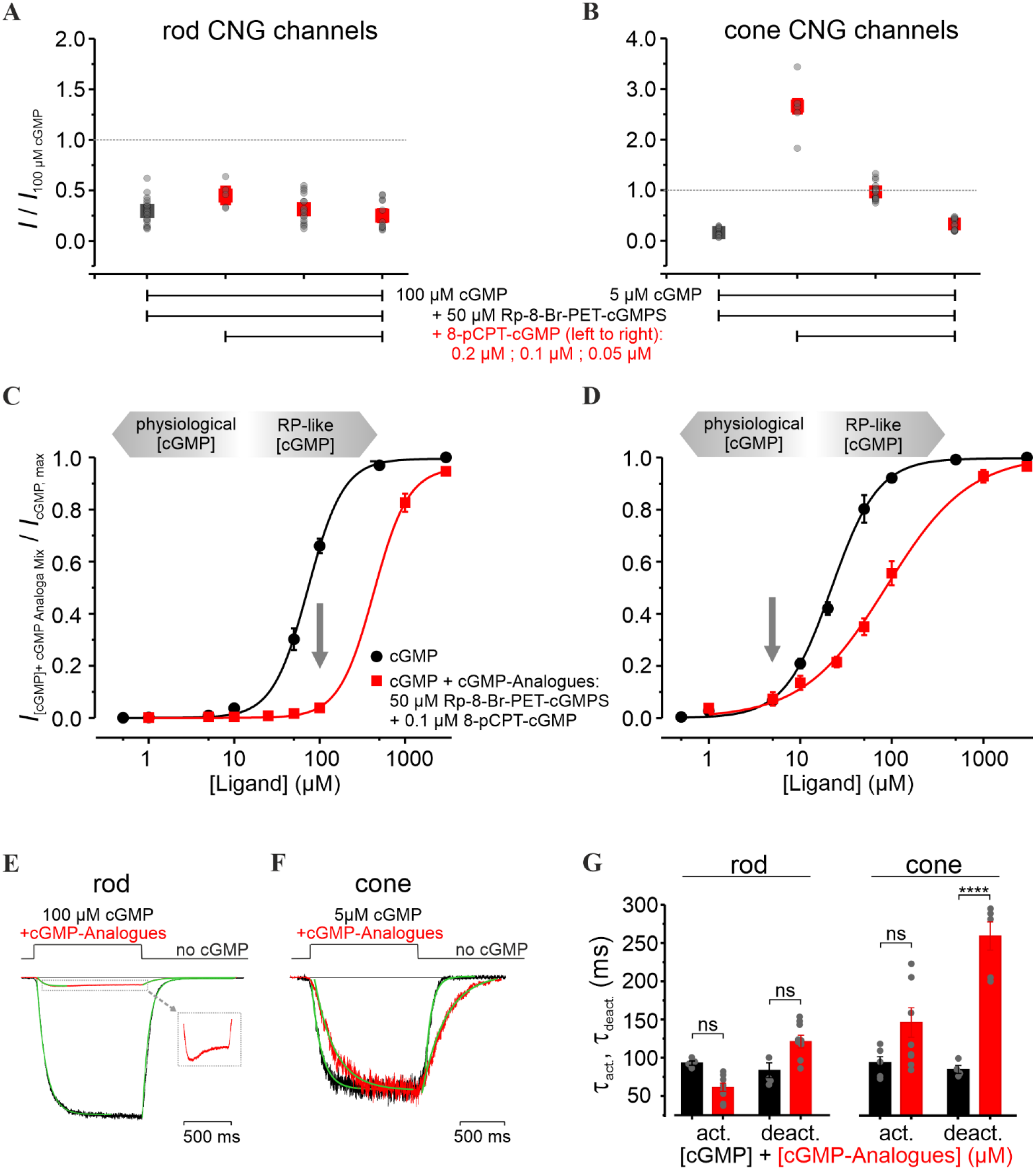
Influence of the combined cGMP-analogues treatment on retinal CNG channels. **A,B**) Relative current amplitudes of rod and cone CNG channels measured under steady-state conditions, in the presence of either cGMP + inhibitor (Rp-8-Br-PET-cGMPS, 50 μM, green) or cGMP + inhibitor (50 μM) + activator (8-pCPT-cGMP, 0.2 μM, 0.1 μM and 0.05 μM, red). The measured currents were related to the activation triggered by 5 μM and 100 μM cGMP for cone and rod channels, respectively. **C,D**) Concentration-activation relationships for rod (left) and cone (right) CNG channels, measured at −35 mV, in the presence of cGMP (black) and cGMP + inhibitor (50 μM) + activator (0.1 μM) (red). The experimental data points, representing means of several measurements were fitted with Eq. (1) (for *EC*_50_, *H* and n see Table S1). cGMP levels <10 μM were considered physiological conditions, cGMP levels >10 μM represent “RP-like”-conditions (grey arrows). **E,F**) Superimposition of representative activation- and deactivation-time courses following a concentration jump from 0 μM cGMP to 100 μM cGMP + 50 μM inhibitor + 0.1 μM activator for rod CNG channels (left) and to 5 μM cGMP + 50 μM inhibitor + 0.1 μM activator for cone (right). The currents triggered by cGMP (black) and by cGMP-analogues co-application (red) were normalized to the level observed in (C,D). The respective time courses were quantified by a monoexponential fit yielding the activation and deactivation time constants (τ_act_ and τ_deact_, respectively, Eq. 3, green curves). The inset in (E) shows the magnified effect of the cGMP-analogues combination on the rod CNG-channel activity (red). **G**) Activation and deactivation time constants (τ_act_, τ_deact_) for rod and cone CNG channels (n = 4-9). The grey symbols represent individual measurements.

Remarkably, the co-application of activator and inhibitor rescued cone-channel activity at physiological cGMP, while inhibiting rod channels at pathological cGMP levels (Fig. 4A,B). The respective concentration-activation relationships confirmed these findings: up to 5 μM cGMP, the cone-channel activity was very similar to the physiological one, while at 100 μM cGMP, the rod-channel activity was considerably reduced (Fig. 4C,D). Moreover, for cone CNG channels, we observed a decrease in the steepness of the concentration-activation relationship (*H* coefficient, Table S1), suggesting that the combined cGMP-analogues treatment reduced the cooperativity during channel gating. This effect was observed only at high cGMP levels, which in RP are not expected to occur in cones (*37*).

Next, we characterized the effect of the cGMP-analogues combination on the channel-gating kinetics. Although the activation level and the activation-time course (τ_act_) of cone CNG channels under this treatment did not change significantly, the deactivation-time course (τ_deact_) was strongly increased (259.0 ± 18.3 ms *vs*. 80.6 ± 4.6 ms in case of cGMP) (Fig. 4F,G). In comparison, rod CNG channels showed almost no change in the gating kinetics (Fig. 4 E,G). Nevertheless, the fast activation phase of the rod channel was followed by an unexpected decrease in the current amplitude to a constant level (Fig 4E, inset). We speculate that this channel behavior mirrors a faster kinetic of the activator effect compared to that of the inhibitor.

### Measuring the effects of cGMP analogues on rod and cone light responses

To study the effects of CNG-channel inhibition and/or activation on photoreceptors maintained in their normal histotypic context, we investigated light-evoked electrical responses in mouse retinal explants, utilizing multi-electrode arrays (MEA). Photoreceptor responses were assessed by measuring the negative deflection in the micro-electroretinogram (μERG), which indicates photoreceptor hyperpolarization and corresponds to the so-called a-wave in a conventional human ERG (*38*) (Fig. 5A upper panel and inset). Responses of retinal ganglion cells (RGC) were measured simultaneously (Fig. 5A lower panel). Since RGCs are third-order neurons, their light-correlated response was delayed compared to that of photoreceptors (*cf*. Fig. 5, right). This temporal correlation between a-waves and the onset of RGC spiking activity was used to determine the timing of the a-wave peak. Moreover, to discriminate between rod and cone photoreceptor specific effects of cGMP-analogues, the recordings were carried out under scotopic (dark adapted, Fig. 5A) and photopic (light adapted, Fig. 5B) conditions, respectively.

**Fig. 5.**
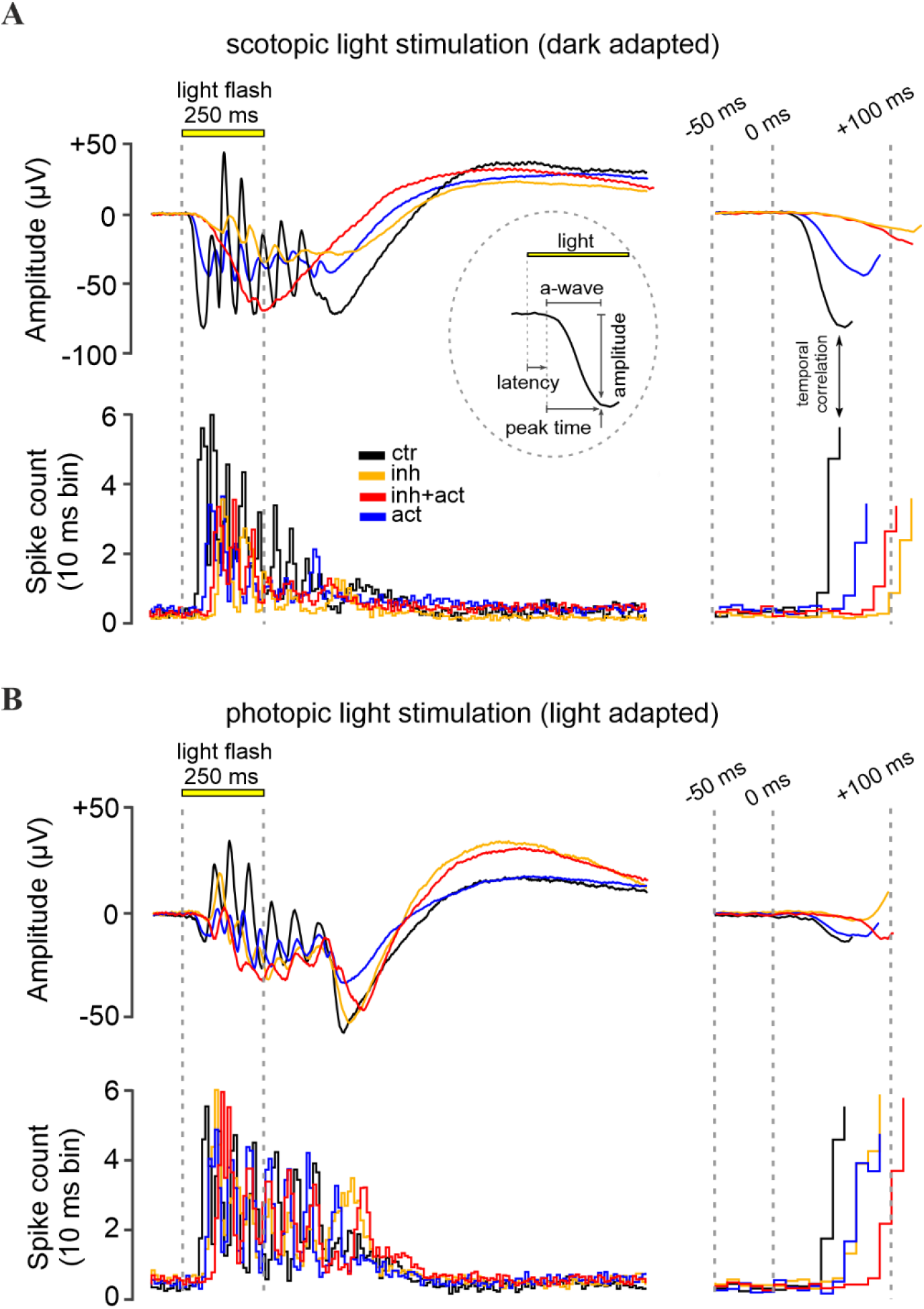
Effects of cGMP analogues on retinal function. Representative light-evoked dark-adapted scotopic (rod) **(A)** and light-adapted photopic (cone) **(B)** micro-electroretinogram (μERG) of the mouse retina. Control recording (black) was compared to retina treated with the CNG-channel inhibitor Rp-8-Br-PET-cGMPS (50 μM, orange), the CNG-channel activator 8-pCPT-cGMP (0.1 μM, blue), or both compounds in combination (red). Displayed below the μERG traces is the correlated retinal ganglion cell spiking activity recorded by multi-electrode arrays (MEA). Data shown at maximum scotopic and photopic light intensity, respectively (see Figs. S3 and S4 for μERG recordings at all light intensities). The left panel illustrates the recorded signals, while the right panel highlights the initial light response features, magnified in time. The initial negative deflection of the μERG, the a-wave, represents the hyperpolarizing photoreceptor response. Subsequent oscillations reflect responses of inner retinal cells. The inset in (A) displays the a-wave evaluation parameters: latency, peak-time, and amplitude.

Under scotopic conditions, below −2.0 log units, rod a-wave amplitudes and RGC spiking activity increased with the light intensity in an almost linear fashion (Fig. 6A, control, black, see also Figs. S3 and S4 for representative traces). At higher light intensities (≥ −2.0 log) retinal responses were larger, yet the response intensities of both a-wave and RGC spiking activity eventually saturated. Since ~ 97 % of the mouse photoreceptors are rods while only ~ 3 % are cones (*39*), the responses recorded within this range were considered as rod-dominant responses. To assess cone responses, the retina was light adapted to saturate rods, so that photopic cone-only responses could be recorded. Due to the low numbers of cones, the photopic a-wave was diminished by ~ 93 %, whereas the RGC spiking responses were comparable to high-light scotopic responses. With increasing light intensities, a slight but continuous increase in the cone a-wave amplitude was observed, while the correlated RGC spike responses appeared to be saturated (Fig. 6B).

**Fig. 6.**
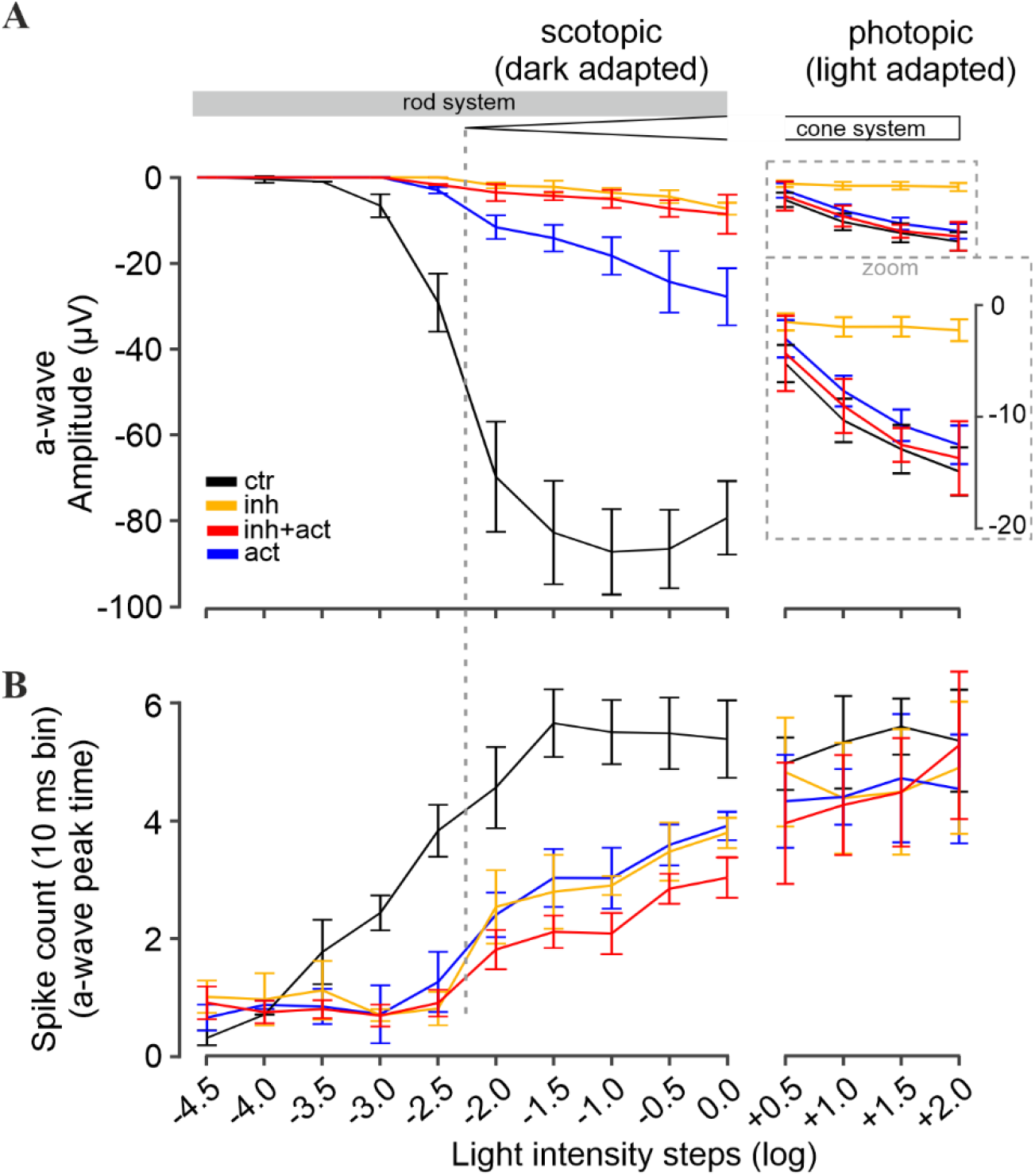
Modulation of retinal light responses by cGMP analogues. Retinal function was measured under scotopic (dark adapted, left panel) and photopic (light adapted, right panel) conditions, reflecting rod and cone function, respectively. **A**) Light-flash mediated photoreceptor responses shown as a-wave peak amplitudes, and (**B**) correlated retinal ganglion cell spike responses shown as counts per 10 ms bin. Experimental conditions: control (black), inhibitor (Rp-8-Br-PET-cGMPS, 50 μM, orange), activator (8-pCPT-cGMP, 0.1 μM, blue) and inhibitor and activator co-application (red, 50 μM and 0.1 μM, respectively). n = 5 retinae for each condition. Data shown averaged from 30 MEA recording electrodes, per condition. See Figs. S3 and S4 for the respective μERG trace and spike data, Tables S2 and S3 for all values and statistical evaluation.

### Rp-8-Br-PET-cGMPS selectively silences rod photoreceptors

Experiments on heterologously-expressed rod and cone CNG channels showed that Rp-8-Br-PET-cGMPS (*i.e*. the “inhibitor”) reduced their activity. Hence, after pre-incubation of photoreceptors with this compound fewer CNG channels remained open that could be closed upon light stimulation, resulting in lower photoreceptor response amplitudes, mimicking light-adaptation (= saturation).

Accordingly, the incubation of the retina with the inhibitor essentially abolished photoreceptor responses under low scotopic light conditions (< −2.0 log) (Fig. 6A, orange). At higher light intensities (≥ −2.0 log) a-wave responses were detectable, even though their amplitudes were strongly reduced (~ 96 %). Yet, RGC spiking activity was only moderately reduced (~ 30 - 40 % drop, Fig. 6B, orange). At higher scotopic stimulation, the a-wave and spiking responses were within the range of photopic cone photoreceptors values. This inhibitor-induced behavior resembled the switch from dark-adapted, scotopic stimulation to light-adapted photopic stimulation in the control situation (Fig. 6A, black): a strong drop of the a-wave amplitude (~ 93%), but no significant alterations of spike responses. Hence, it appears that the inhibitor selectively silenced rod responses. In other words, the inhibitor treatment and the subsequent closure of CNG channels emulated a light-dependent saturation of rods, allowing for cone recordings during scotopic stimulation.

In addition, under light adapted photopic stimulation (≥ +0.5 log), a response-dampening effect of the inhibitor on cone photoreceptors was observed (Fig. 6A). In control recordings, the a-wave amplitudes increased linearly with increasing light intensities. In contrast, in the presence of the inhibitor, the cone a-wave amplitudes showed only minimal increases. On the other hand, the RGC spiking activity remained nearly at control levels.

Overall, the results indicate that the inhibitor silenced rod photoreceptor light responses selectively, both in terms of a-wave amplitudes and RGC spiking activity. Importantly, cone a-waves were only dampened by the inhibitor, with cone-dependent RGC activity remaining at control levels.

### 8-pCPT-cGMP counteracts the inhibitor effects in cones but not in rods

Next, we investigated the potential of 8-pCPT-cGMP (*i.e*. the “activator”) to counteract the response dampening effect of the inhibitor on photoreceptors operating at physiological cGMP levels. In experiments with heterologously-expressed rod and cone CNG channels the activator behaved as potent agonist and thus created a situation equivalent to dark-adaptation. Interestingly, when the mouse retina was incubated with the activator and inhibitor mixture, the activator countered the effect of the inhibitor with a significant preferential for cones. The photopic a-wave as well as the spike responses (Fig. 6, red) were within the control range, with no significant difference. In the scotopic range, however, the rod response dampening effect of the inhibitor remained dominant, even though, the activator produced a minor rescue effect also here: The rod a-wave amplitude increased, and the response was detectable at −2.5 log units, *i.e*. 0.5 log units earlier in comparison to inhibitor alone (at −2.0 log). While this improvement of a-wave amplitudes was significant, it was still about ~ 90 % lower than control (> 2.5 log). On the other hand, the spiking responses of RGCs were similar to control, especially in the photopic range, again suggesting a cone selective rescue in the presence of the activator and inhibitor mixture.

The incubation of the retina with the activator alone revealed no effect on cones (Fig. 6, ≥ + 0.5 log), but instead produced a marked shift in the sensitivity of rods. Under low scotopic light stimulation rod responses were essentially absent (< −2.0 log range of pure rod responses, Fig. 6A, blue). At higher light intensities (≥ −2.0 log), the a-wave amplitudes of the photoreceptors were reduced by 60 - 80 % (Fig. 6A), while light-mediated RGC spike responses dropped by only 35 - 47 % (Fig. 6B). Moreover, in contrast to control, the response intensities (a-wave and RGC spikes) increased linearly with rising light intensities, suggesting that the rods were not responding at their full capacity (*i.e*., rods were not saturated). Taken together, our results indicated an activator-mediated shift in rod sensitivity such that higher light intensities were required to elicit responses.

### cGMP analogues modulate the kinetics of photoreceptor responses

We proceeded further by analyzing the μERG data in the temporal domain, evaluating the a-wave slope (μV per 20 ms) as an indirect measure for the light-intensity and time-dependent closure of CNG channels. During scotopic stimulation, the control a-wave slope followed the trend of the a-wave amplitudes: they increased linearly with rising light intensities (< −2.0 log, *i.e*., rod only range, from 0 to −2.20 ± 0.42 μV/20ms) and eventually reached saturation at −2.57 ± 0.28 μV/20 ms for light intensities above −2.0 log (Fig. 7 left, control, black). In contrast to control, the inhibitor produced a marked (up to 90 %) and significant decrease in the a-wave slope (orange). Since the inhibitor essentially silenced the rod system (*cf*. Fig. 6), the responses obtained within the scotopic range likely relate to the cone system. Remarkably, the co-application of inhibitor & activator (Fig. 7, left, red), did not improve the a-wave slope when compared to inhibitor only. On the other hand, the activator treatment alone (blue), decreased the a-wave slope by ~ 85 % in comparison to control, but this value was still significantly higher than those for the inhibitor (orange) and the inhibitor & activator mixture (red).

**Fig. 7.**
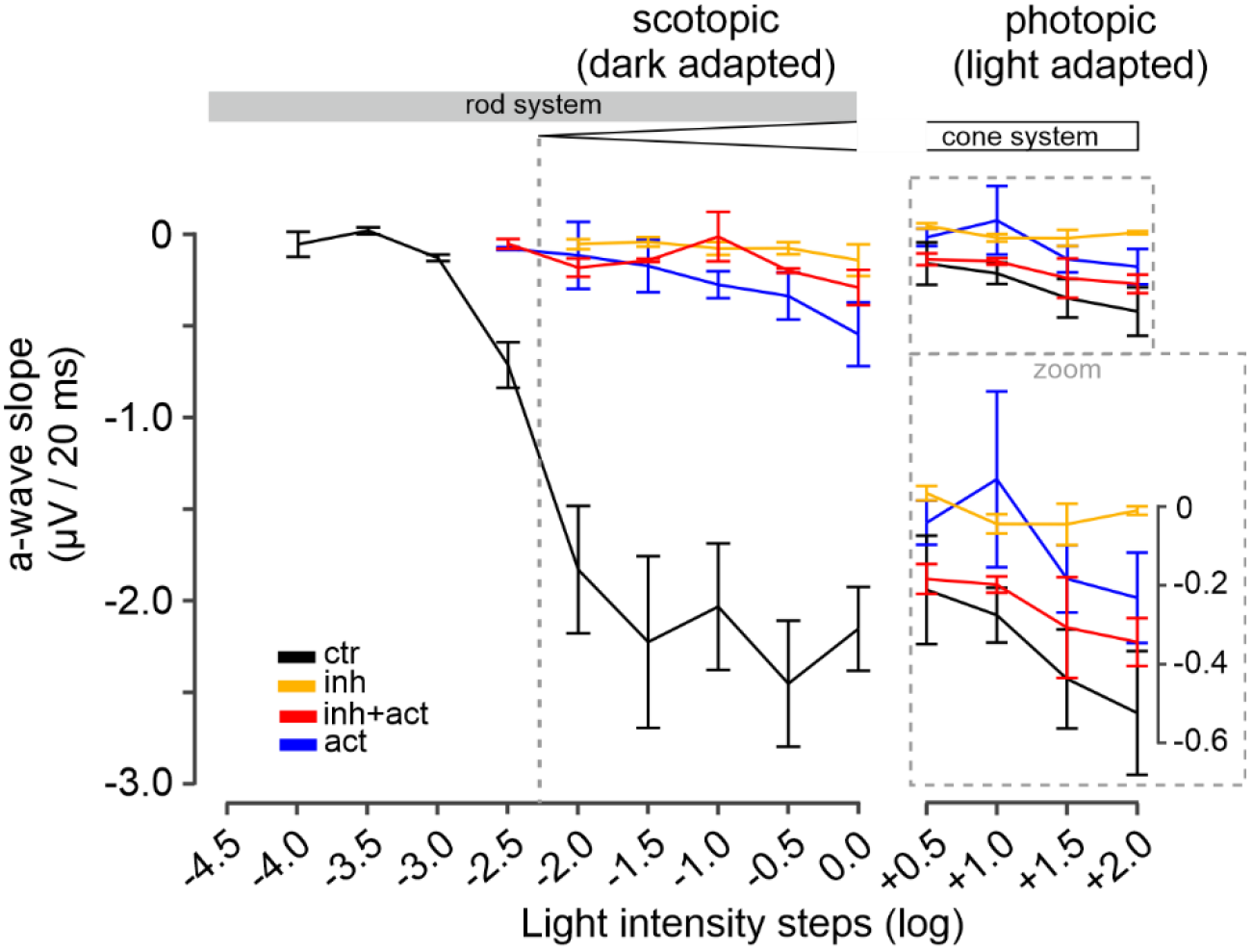
Modulation of photoreceptor-response kinetics by cGMP analogues. Illustration of the a-wave slope calculated as a-wave amplitude deflection in μV per 20 ms. Scotopic (dark adapted, rod function) light conditions shown in left panel, photopic (light adapted, cone function). Experimental conditions: control (black), inhibitor (Rp-8-Br-PET-cGMPS, 50 μM, orange), activator (8-pCPT-cGMP, 0.1 μM, blue) and inhibitor & activator co-application (red, 50 μM and 0.1 μM, respectively). n = 5 retinae for each condition. Data shown are averages of 30 MEA recording electrodes, per condition. See Figs. S3 and S4 for the respective μERG trace and spike data, Tables S4 and S5 for all values and statistical evaluation.

Under photopic conditions, the control cone a-wave slopes showed a relatively minor increase with light intensity, with no sign of saturation, ranging at an average level of −0.36 ± 0.14 μV/20 ms (Fig. 7 right, control, black). For the inhibitor (orange), along with the reduced a-wave amplitude, also a prominent decrease in the a-wave slope was measured (on average 90 %). In contrast, the mixture of inhibitor & activator (Fig. 7, right, red) produced a 80 - 90 % recovery of the a-wave slope in comparison to control (see Figs. S4 and S5 for values and statistical evaluation). The activator alone, however, decreased the a-wave slope by ~ 60 % (averaged) in comparison to control (blue). Overall, the analysis of the photoreceptor-response kinetics supported a cone-selective modulation for the activator and a rod-selective modulation for the inhibitor.

## DISCUSSION

### Targeting the rod photoreceptors

Here, we present a novel approach to selectively inhibit rod CNG channels while preserving cone functionality. We employed patch-clamp recordings to characterize direct drug effects on channel activity, and then used the newly developed μERG technique to detect treatment-induced voltage changes in rod and cone photoreceptors maintained in their natural histotypic context. The latter were correlated with retinal ganglion cell spiking activity, allowing to determine the precise maxima of photoreceptor hyperpolarization. Together, these complementary methods enabled a comprehensive characterization of our novel CNG channel targeting approach, also illustrating the potential to selectively alter rod and cone photoreceptor visual responses.

Despite our best efforts, we could not identify selective inhibitors for rod CNG channels. Even Rp-cGMPS, a known PKG inhibitor previously reported to have opposite effects on rod and olfactory CNG channels (*36, 40*), showed no clear preference for any of the retinal CNG-channel isoforms. All tested compounds either shifted the activation range of both rod and cone CNG channels to higher ligand concentrations or had no effect at all. Under RP-like cGMP conditions, the co-application of cGMP analogues with antagonistic functional characteristics lead to the inhibition of rods and cones, combined with a selective activation of cone CNG channels. Thus, primary survival of rods due to CNG-channel inhibition should promote secondary cone survival at functional level.

This selective modulation was confirmed also on intact rod and cone photoreceptors operating at natural cGMP levels. While on heterologously-expressed CNG channels the inhibitor alone displayed only a weak selectivity for rod channels, in the retina the inhibitor essentially silenced rod activity. The combined cGMP-analogues treatment helped recovering the cone responses from the inhibitor’s effect, raising additional questions about the modulation mechanism of the activator. The activator showed a strong concentration-dependent cone CNG-channel selectivity and maintained the normal function of the cone photoreceptors. Interestingly, the activator, under scotopic condition prevented a response saturation of rods – even at higher light intensities – indicating a modulation of rod-response kinetics. Our data suggest that under illumination, in the presence of the activator, when cGMP is hydrolyzed, the CNG channels close only very briefly, just as long as it takes the activator to reopen them. In other words, the activator forces the photoreceptor into an artificial “dark-adaptation”. To illustrate the effects of the different compounds on the response amplitudes of individual rod and cone photoreceptors, we compiled a model that also shows their likely effects on the resting membrane potential (Fig. 8).

**Fig. 8.**
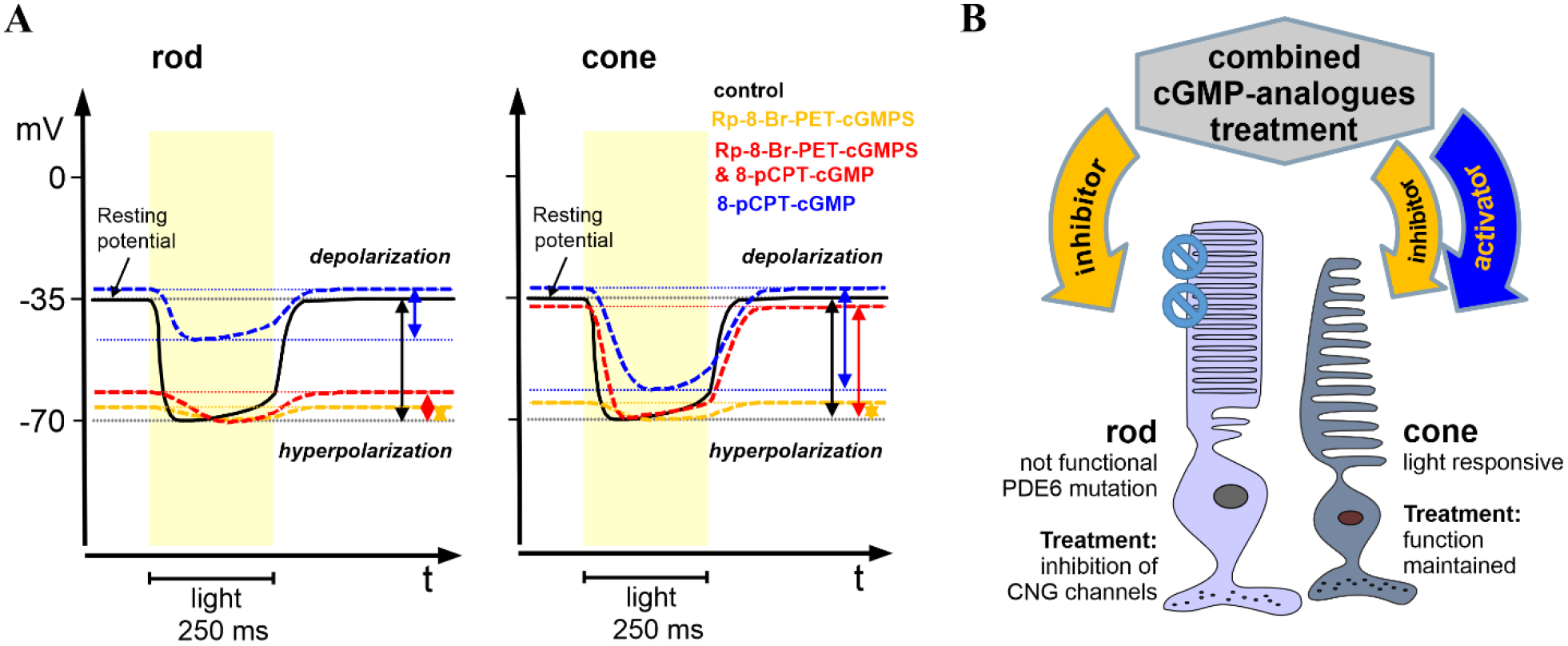
Effect of the combined cGMP-analogues treatment on photoreceptors. **A)** Model for the effects of cGMP analogues on rod and cone photoreceptors from wt-retina. Idealized representation of photoreceptor membrane potentials in different experimental conditions. Under control conditions (black traces), in the dark, rod (left) and cone (right) photoreceptors maintain a resting potential of −35 mV and respond to light with a hyperpolarization to approx. −70 mV (black arrows) (*5*). The inhibitor (Rp-8-Br-PET-cGMPS, orange) reduces the influx of positive charges (Na^+^/Ca^2+^) in both rods and cones, shifting the resting potential to more negative values. Since most CNG-channels are already closed by the inhibitor (*i.e*. mimicking constant light), the light-induced hyperpolarization response is strongly decreased compared to control (orange arrows). In cone photoreceptors, the activator (8-pCPT-cGMP, blue) increases the influx of cations, raising the resting potential to more positive values (*i.e*. mimicking constant darkness). In light, this leads to decreased cone response amplitudes compared to control (blue arrows). The combination of inhibitor & activator (Rp-8-Br-PET-cGMPS + 8-pCPT-cGMP; red) partially offsets the effect of channel inhibition, slightly increasing cone response amplitudes (red arrows). In rod photoreceptors, the activator alone (blue) decreases light-induced rod response amplitudes. Moreover, the combined application of inhibitor and activator in rods produces an effect on response amplitudes similar to the inhibitor alone. Note that for the drug treatments the photoreceptor voltages are only rough approximations, based on the relative differences between response amplitudes recorded for the different experimental conditions. **B**) Strategy of selective CNG-channel isoform modulation. Selective inhibition of rod CNG channels under RP-like cGMP conditions was achieved by co-application of a weak-selective rod CNG-channel inhibitor and of a cone CNG-channel selective activator.

### The combined cGMP-analogues treatment modulates kinetics of photoreceptor responses

Given the highly dynamic nature of our surrounding, the ability of the human eye to promptly register and identify changes is critical for survival. Therefore, for an efficient RP treatment, the cGMP analogues should elicit their effect without influencing the responsiveness of photoreceptors. Although the inhibitor-activator treatment was intended to target the rod photoreceptors only, we observed a slower deactivation kinetics of cone CNG channels. The reason for this might be slower ligand unbinding, or slower conformational changes within the channel protein upon unbinding, or a combination of both. Surprisingly, at the photoreceptor level, with photopic stimulation and the combined inhibitor-activator treatment, the slope of cone-cell hyperpolarization was barely changed. This suggests, as also previously discussed (*41*), that the a-wave slope cannot be simply correlated to the kinetics of CNG-channel deactivation only, as it represents a reflection of the photocurrent, *i.e*. the end product of the phototransduction cascade.

Although the kinetics of the triggered membrane potential change was unaffected, under the inhibitor-activator treatment, we observed also a delayed light response of cone photoreceptors. Whether this delay is physiologically relevant for the light-adaptation mechanism or visual performance in general could be assessed in future *in-vivo* experiments on RP-animal models. Still, even if adverse changes in kinetics were to be observed, the benefits of preserving vision in RP patients would likely outweigh such side-effects. In addition, confocal patch-fluorometry with fluorescently-labeled ligands could be employed to study the binding of cGMP-analogues to retinal CNG channels (*42*). This could help us to better understand the way of action of these cGMP analogues and possibly to identify and address the changes in the photoreceptor’s responsiveness.

### Therapeutic implications for the combined cGMP-analogues treatment in retinal degenerative diseases

We have shown that the CNG-channel inhibitor Rp-8-Br-PET-cGMPS, previously reported to delay photoreceptor degeneration in several RP mice models (*27*), when applied alone, was not rod selective, but also reduced the light-induced responses in cones. To our knowledge, the strategy of combining ligands with agonistic and antagonistic behaviors to achieve channel selectivity has never been described before and it is tempting to think of it in terms of a personalized and balanced-drug treatment. The tuning of the cGMP-analogues composition and their respective concentration to selectively control the diseased photoreceptors is a multilevel approach. It requires a basic understanding of their direct effect on cGMP-dependent channels (1), on intact photoreceptors with naturally-balanced cGMP levels (2) and on diseased retinal tissue (3). Our study covered the first two levels of this approach. Future *in vivo* studies on different RP-animal models may further elucidate the effect of this treatment and its influence on animal’s visual performance.

Nevertheless, one has to be aware of possible challenges arising due to the genetic heterogeneity of different RP forms, different cGMP levels in the diseased cells and different kinetics of the disease development (*17, 43*). Still to be determined is the effect of this cGMP-analogues combination also on other CNG-channel isoforms (*e.g*. olfactory CNG channels). Despite the multitude of studies regarding the CNG channel-gating mechanism, there are still many open questions with respect to ligand selectivity among channel isoforms (*5*). Important steps in this direction were made in 2017 when the first full-length structure of a cGMP-bound open-state CNGA1 channel from *C. elegans* was published and later, from 2020 to 2022, when several rod and cone CNG-channel structures were reported (*44–48*). This sudden wealth of structural information will likely trigger future cGMP-binding studies and discovery of novel ligand-protein interactions with determinant roles in ligand selectivity. However, it will still be challenging to design selective cGMP analogues since the structure of the ligand-binding domain is highly conserved within the CNG channel family, making the strategy to use the combined cGMP-analogues treatment even more attractive.

Future studies should also address the effect of such combination treatments also on other potential cellular targets containing cGMP-binding domains, such as cyclic nucleotide phosphodiesterases (PDEs), or hyperpolarization-activated, cyclic nucleotide-modulated (HCN) channels. Another key aspect for the success of our combination treatment will be the kinetics of delivery for the two different cGMP analogues to the photoreceptors. Here, a suitable delivery approach must ensure that both compounds can exert their effects within same time frame (*49*).

Our data could also be of strong interest for the treatment of *achromatopsia*, a rare autosomal recessive cone disorder where the rod-mediated vision remains largely unaffected (*50*). Patients suffering from *achromatopsia* experience strong photophobia due to saturation of rod photoreceptors in day light (*51*). Here, a potential therapy could aim at decreasing the sensitivity of rod photoreceptors, such as was shown in this study for the activator treatment. Overall, the implications of our findings may go beyond the treatment of retinal diseases, and could also include the selective regulation of rod and/or cone photoreceptor sensitivity, to improve night or daylight vision, respectively. Furthermore, the combination of an inhibitor-activator treatment may not be limited to CNG channels but may be applicable also to other disease conditions, where targeting the proteins of interest was up to now a challenge.

## MATERIALS AND METHODS

### Molecular biology and heterologous expression of photoreceptor CNG channels in *Xenopus laevis* oocytes

For heterologous expression in *Xenopus laevis* oocytes, bovine CNGA1 (NM_ 174278.2) and CNGB1a subunits (NM_181019.2) from rod photoreceptors and human CNGA3 (NM_001298.2) and CNGB3 subunits (NM_019098.4) from cone photoreceptors, were subcloned into the pGEMHE vector (*52*). The cDNA encoding for the human cone CNGB3 subunit was kindly provided by M. Varnum (Washington State University, Department of Veterinary and Comparative Anatomy, Pharmacology and Physiology, USA) and for the bovine CNGB1a subunit by W. Zagotta (University of Washington, Department of Physiology and Biophysics, USA).

Oocytes were surgically removed from adult females of *Xenopus laevis* under anesthesia with 0.1% tricaine, pH = 7.1 (MS-222, Parmaq, UK) and they were incubated for 105 min in Ca^2+^-free solution (in mM: 82.5 NaCl, 2 KCl, 1 MgCl2, and 5 Hepes, pH 7.4) containing 3 mg/mL collagenase A (Roche Diagnostics, Germany). Oocytes of stage IV and V were manually defolliculated and each oocyte was injected with 50-100 ng of mRNA encoding for the respective photoreceptor CNG channels. The vitelline membrane was manually removed from the oocyte before the electrophysiological experiments. For efficient generation of heterotetrameric channels, the ratio of CNGA3 mRNA to CNGB3 mRNA was 1:2.5 (*7*) and of CNGA1 mRNA to CNGB1a mRNA was 1:4 (*33, 53*). After the mRNA-injection the oocytes were kept at 18°C for 2 to 7 days in a solution containing (in mM) 84 NaCl, 1 KCl, 2.4 NaHCO_3_, 0.82 MgSO_4_, 0.41 CaCl_2_, 0.33 Ca(NO_3_)_2_, 7.5 Tris, pH 7.4.

### Electrophysiology

Patch-clamp recordings were performed on inside-out patches from *Xenopus laevis* oocytes expressing heterotetrameric rod and cone CNG channels. Recordings were made at room temperature using an Axopatch 200B patch-clamp amplifier (Axon Instruments, Foster City, CA). Electrophysiology was controlled by the PatchMaster-software (HEKA Elektronik Dr. Schulze GmbH, Lambrecht, Germany). The sampling rate was 5 kHz and the filter implemented in the amplifier was set to 2 kHz. From a holding potential of 0 mV, currents were elicited by voltage steps to −100 mV, then to +100 mV, and back to 0 mV. When mentioned, also voltage steps to −35 mV, the physiological voltage under dark conditions in photoreceptors, were applied.

Intracellular and extracellular solutions contained 140 mM NaCl, 5 mM KCl, 1 mM EGTA, and 10 mM HEPES (pH 7.4). The cyclic nucleotides, cAMP (Merck KGaA, Darmstadt, Germany), cGMP and cGMP analogues (Biolog LSI GmbH & Co. KG, Bremen, Germany) were added to intracellular solutions as indicated. The concentrations of the respective dilutions were verified by UV-spectroscopy (Thermo NanoDrop 2000c, Germany). The L-cis-diltiazem solutions (100 μM, Abcam, Germany) were prepared from stock solutions (10 mM) shortly before the measurements. The patch pipettes were pulled from borosilicate glass tubing (outer diameter 2.0 mm, inner diameter 1.0 mm; Hilgenberg GmbH, Germany). The initial resistance of the solution-filled pipettes was 0.6-1.4 MΩ. The cGMP-containing solutions were administered via a multi-barrel application system to the cytosolic face of the patch. For studying the activation and deactivation kinetics of CNG channels, we performed fast jumps between different ligand concentrations by means of a double-barreled θ-glass pipette mounted on a piezo-driven device, which was controlled by the computer. For measuring the time courses, the recording rate was 20 Hz. The effective switch time of the solution exchange, determined with an open patch pipette and different solutions in the barrels, was negligible compared to the time courses of channels activation and deactivation (*54*).

### cGMP analogues

All cGMP analogues (Rp-cGMPS, Rp-8-Br-(2-N)ET-cGMPS (DF156), Rp-(2-N)ET-cGMPS (DF246), and Rp-ß-1,N2-Ac-Br-cGMPS (DF235)) were diluted from stock solutions according to the technical details provided by the manufacturer (Mireca Medicines GmbH, Tübingen and Biolog GmbH & Co. KG, Bremen, Germany). cGMP analogues (see Fig. 1) were prepared according to previously reported methods (*27*) and US20190292214 (“New equatorially modified polymer linked multimers of guanosine-3,5-cyclic monophosphates”).

### Steady-state concentration-activation relationships

Each patch was first exposed to a solution containing no cGMP and then to a solution containing a saturating cGMP concentration. After subtracting the current in the absence of cGMP, the current response for each ligand concentration was normalized to the saturating current. The respective concentration-activation relationships were fitted with a Hill equation:

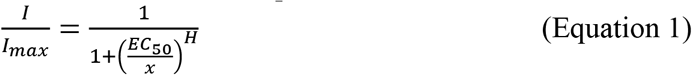

where *I* is the current amplitude, *I*max is the maximum current induced by a saturating ligand concentration, *x* is the ligand concentration, *EC*_50_ is the ligand concentration of half maximum effect, and *H* is the Hill coefficient. The concentration-activation relationship describing the inhibitory effect of Rp-8-Br-PET-cGMPS was best fitted by a double Hill equation:

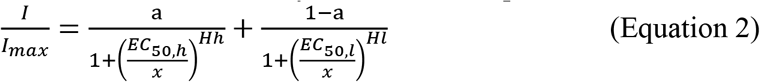

where *I* is the current amplitude, *I*_max_ is the maximum current induced by a saturating ligand concentration, *x* is the ligand concentration, *EC*_50,h_ and *EC*_50,l_ are the ligand concentrations of half maximum effect for the high- and the low-affinity component and *H*_h_ and *H*_l_ are the respective Hill coefficients. *a* and (1-*a*) represent the amplitudes of the high- and the low-affinity components. The activation and deactivation time courses were determined by fitting the respective current traces with single exponentials:

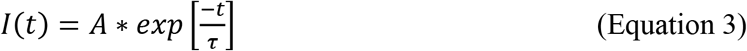

where *A* is the amplitude, *t* the time, and τ the time constant for either activation or deactivation. To be able to differentiate between the effects of the cGMP analogues on the photoreceptor CNG channels, the respective ligands were initially tested on channels that were in a similar activation state (*e.g*. ~80% activation in the presence of 20 μM cGMP for cone and 100 μM cGMP for rod channels). In this way, we made sure that the observed effects were not the result of cGMP-analogue-dependency on the functional state of the channel. For all experiments, we used 50 μM of the cGMP analogues because this concentration induced a strong neuroprotective effect, with no side effects in RP-animal models (*27*).

### Animals

The procedures regarding the *Xenopus laevis* frogs had approval from the authorized animal ethics committee of the Friedrich Schiller University Jena. The respective protocols were performed in accordance with the approved guidelines. Extreme efforts were made to reduce the stress and to keep the number of frogs to a minimum.

In this study, adult postnatal day (P) 25-30 C57BL/6J mice (*n* = 5) were used. Animals were housed under standard 12 hours white cyclic lighting, had free access to food, water and playground, and were used irrespective of gender. Animal protocols compliant with §4 of the German law of animal protection were reviewed and approved by the Tübingen University committee on animal protection. Moreover, experiments were performed in accordance with the ARVO statement for the use of animals in ophthalmic and visual research. All efforts were made to minimize the number of mice used and their suffering.

### Tissue preparation

The mice were kept overnight in a ventilated light-tight box for dark-adaption. Subsequent procedures were performed under dim red light. The eyes of the mice were enucleated, and the retina was isolated in extracellular solution containing (in mM): 125 NaCl, 26 NaHCO_3_, 2.5 KCl, 2 CaCl_2_, 1 MgCl_2_, 1.25 NaH_2_PO_4_, and 20 glucose, and was maintained at pH 7.4 using carboxygen perfusion (95/5% O_2_/CO_2_). All chemicals were obtained from Merck KGaA (Darmstadt, Germany). For MEA-recording the retinas were placed in the MEA-recording chamber (GC side down)(*55*). During the recordings, the tissue was perfused with carboxygenated medium at 32°C. The chamber perfusion rate was adjusted to 2 ml/min.

### Retinal recordings

Electrophysiological recordings were performed to access the light dependent retinal responses (photoreceptors μERG and RGCs spikes) by means of multi-electrode array system (MEA, USB-MEA60-Up-BC-System-E from Multi Channel Systems) equipped with MEA 200/30iR-ITO-pr. The recordings were performed at 25000 Hz sampling rate collecting unfiltered raw data. The trigger-synchronized operation of the light stimulation (LEDD1B T-Cube, Thorlabs) and MEA-recording were controlled by the recording protocol set within the MCRack software (v 4.6.2, MCS) and the digital I/O – box (MCS). The light stimulation (white light LED, 2350 mW, MCWHD3, Thorlabs) was applied from beneath the transparent glass MEA guided by optic fiber and optics. A spectrometer USB4000-UV-VIS-ES (Ocean Optics, USA) was employed to calibrate the intensity of the applied light stimulation protocol. To discriminate the rod and cone photoreceptor responses, a light stimulation protocol for *ex-vivo* MEA recordings was established following the standard *in-vivo* ERG protocol (*56*). For scotopic recordings (rod responses), the animals were dark-adapted (12 hours), while for the photopic recordings (cone responses) the retinal explants were light adapted on the MEA (5 min at intensity of 4.20E+13 photons/cm^2^/sec), subsequent to scotopic stimulation. The parameters of the light stimuli were set as: Light flash 250 ms long (3 repetition per light intensity, 20 s interval). Scotopic: 10 stimuli in 0.5 log steps from −4.5 to 0.0 (1.33E+09 - 4.20E+13 photons/cm^2^/sec). Photopic (background light: 4.20E+13 photons/cm^2^/sec): four stimuli in 0.5 log steps from +0.5 to +2.0 (1.33E+14 - 4.20E+15 photons /cm^2^/sec). The cGMP-analogues were bath-applied and after a 45 min incubation light-evoked μERGs as well as the RGC spikes were recorded from the same retinal patch.

### Data analysis & statistics

The statistical analysis of the data from the heterologously-expressed CNG channels was performed using the two-tailed unpaired Student *t*-test. Experimental data are given as mean ± SEM. Analysis of the experimental data was done with the OriginPro 2016G software (OriginLab Corporation, Northampton, USA).

MEA-recording raw data files were filtered employing the Butterworth 2nd order (MC-Rack, Multi-channel systems) to extract the μERG (field potentials: bandpass 0.01 – 100 Hz) and spikes (high pass 200 Hz). The filtered data were converted to *.hdf files by MC DataManager (v1.6.1.0) for further data processing in MATLAB (*55*). Shown traces data are average of 30 MEA recording electrodes, per condition. n = 5 retinae were recorded per condition. Statistical significance was estimated by one-way ANOVA followed by the Dunnett’s multiple comparisons. Experimental data are given as mean ± SEM.

The a-wave slope was calculated as the quotient of the a-wave deflection in μV per 20 ms period (last 20 ms of the a-wave amplitude; between starting point of the a-wave (response latency) and the ending at the a-wave (peak time), *cf*. Fig. 5, circle).

Figures were prepared using CorelDraw^®^ 2019 (Corel, Ottawa, Canada) and Inkscape (inkscape.org).

## Supplementary Materials

### Supplementary Figures

**Fig. S1.**
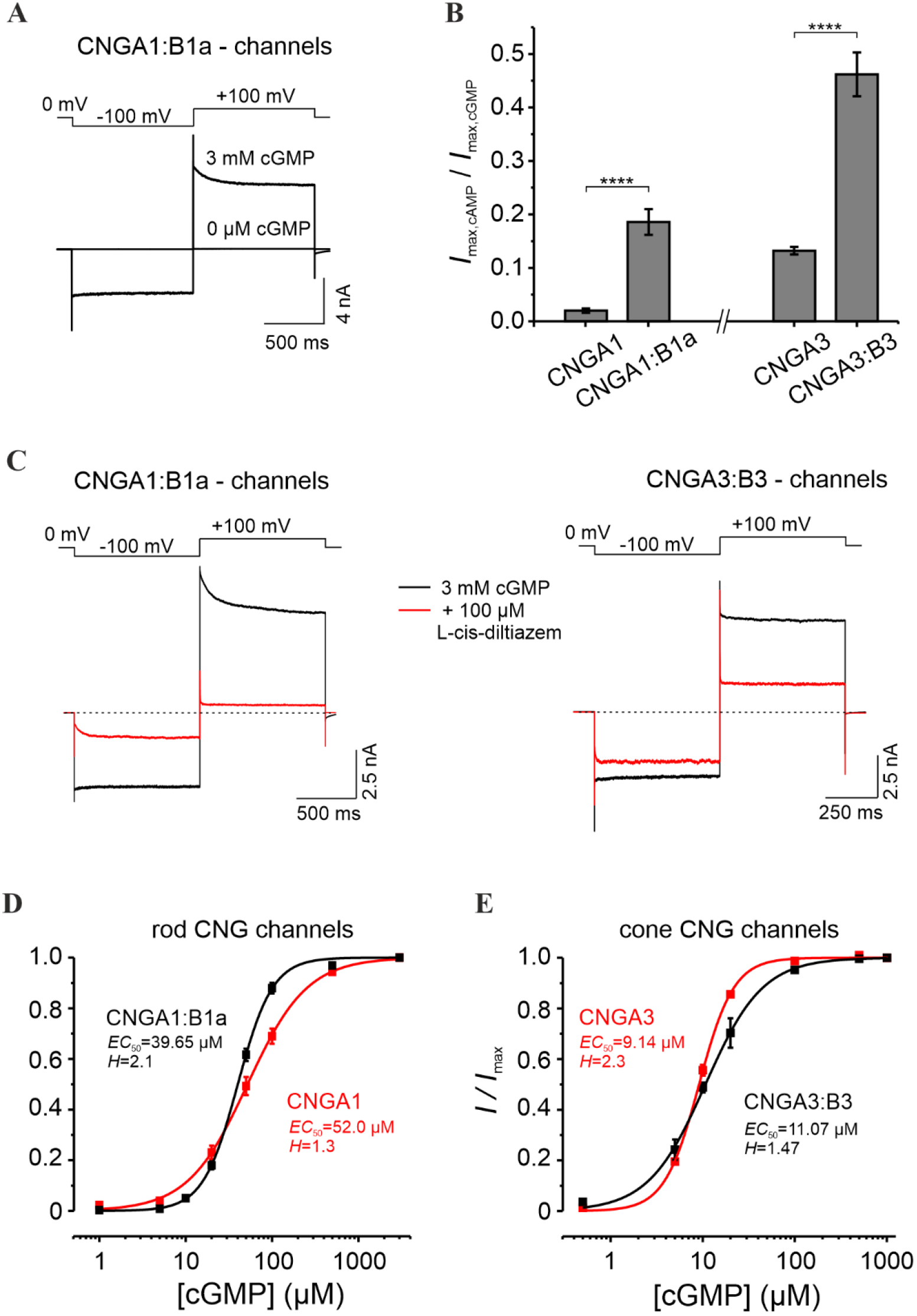
Electrophysiological characterization of rod and cone heterotetrameric CNG channels expressed in *Xenopus laevis* oocytes. **A)** Representative current traces for rod CNGA1:B1a channels in either the absence or the presence of saturating cGMP. The current traces were elicited by voltage steps from a holding potential of 0 mV to −100 mV, +100 mV and 0 mV. Leak currents in the absence of cGMP were subtracted from all recordings. **B)***I*_cAMP_/*I*_cGMP_ at +100 mV was: 0.02 ± 0.004 for CNGA1 (n = 18), 0.19 ± 0.02 for CNGA1:B1a (n = 16), 0.13 ± 0.007 for CNGA3 (n = 20) and 0.46 ± 0.04 for CNGA3:B3 channels (n = 19). **C)** Representative cGMP-activated currents from CNGA1:B1a- and CNGA3:B3-channels with (red) or without (black) 100 μM L-cis-diltiazem. The voltage protocol is shown at the top. At +100 mV, L-cis-diltiazem blocked ~ 90.5 % of rod- and ~ 63.4 % of cone-CNG channel activity triggered by 3 mM cGMP (n = 4). **D,E**) cGMP-dependent concentration-activation relationships for heterotetrameric rod CNGA1:B1a (**D**) and cone CNGA3:B3 (**E**) channels. The experimental data points, each representing the mean (± SEM) of 5 to 14 measurements, were fitted with Eq. (1).

**Fig. S2.**
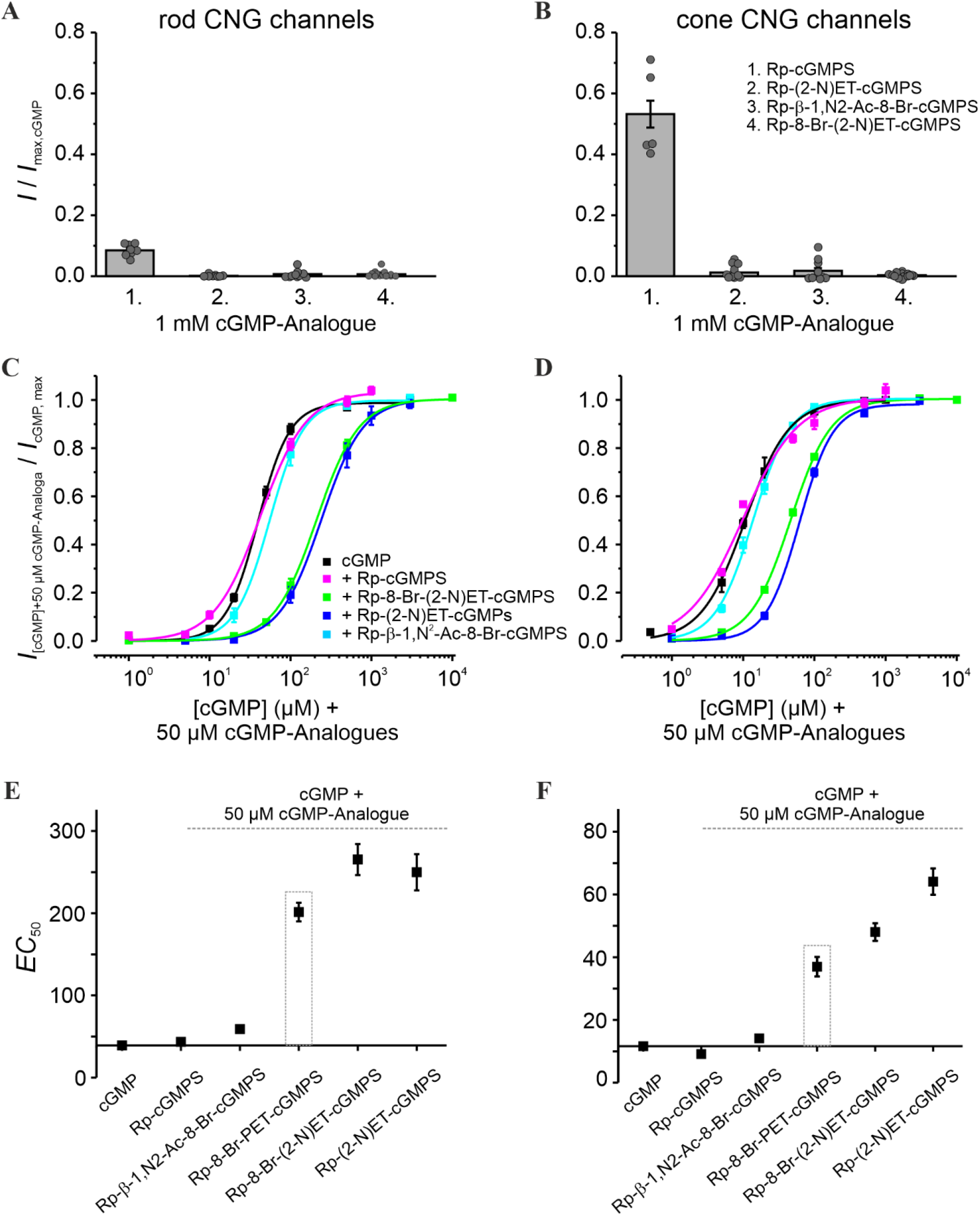
Influence of Rp-modified cGMP analogues on cGMP-induced activation of rod and cone CNG channels. **A,B)** Efficacy of Rp-8-Br-(2-N)ET-cGMPS, Rp-β-1,N^2^-Ac-Br-cGMPS, Rp-cGMPS and Rp-(2-N)ET-cGMPS when activating rod (**A**) and cone CNG channels (**B**). The currents measured at +100 mV, in the presence of the respective cGMP-analogues (1 mM), were related to the maximal cGMP-current. **C,D)** Concentration-activation relationships for rod and cone CNG channels, obtained in the presence of cGMP and the respective cGMP analogues. The currents triggered by the cGMP analogues were normalized with respect to the ones obtained at saturating cGMP. The experimental data points, representing means of several measurements, were fitted with Eq. (1) (for *EC*_50_, *H* and n see Table S1). **E,F)***EC*_50_ values for rod and cone CNG channels. The dotted box underlines the values obtained for Rp-8-Br-PET-cGMPS.

**Fig. S3.**
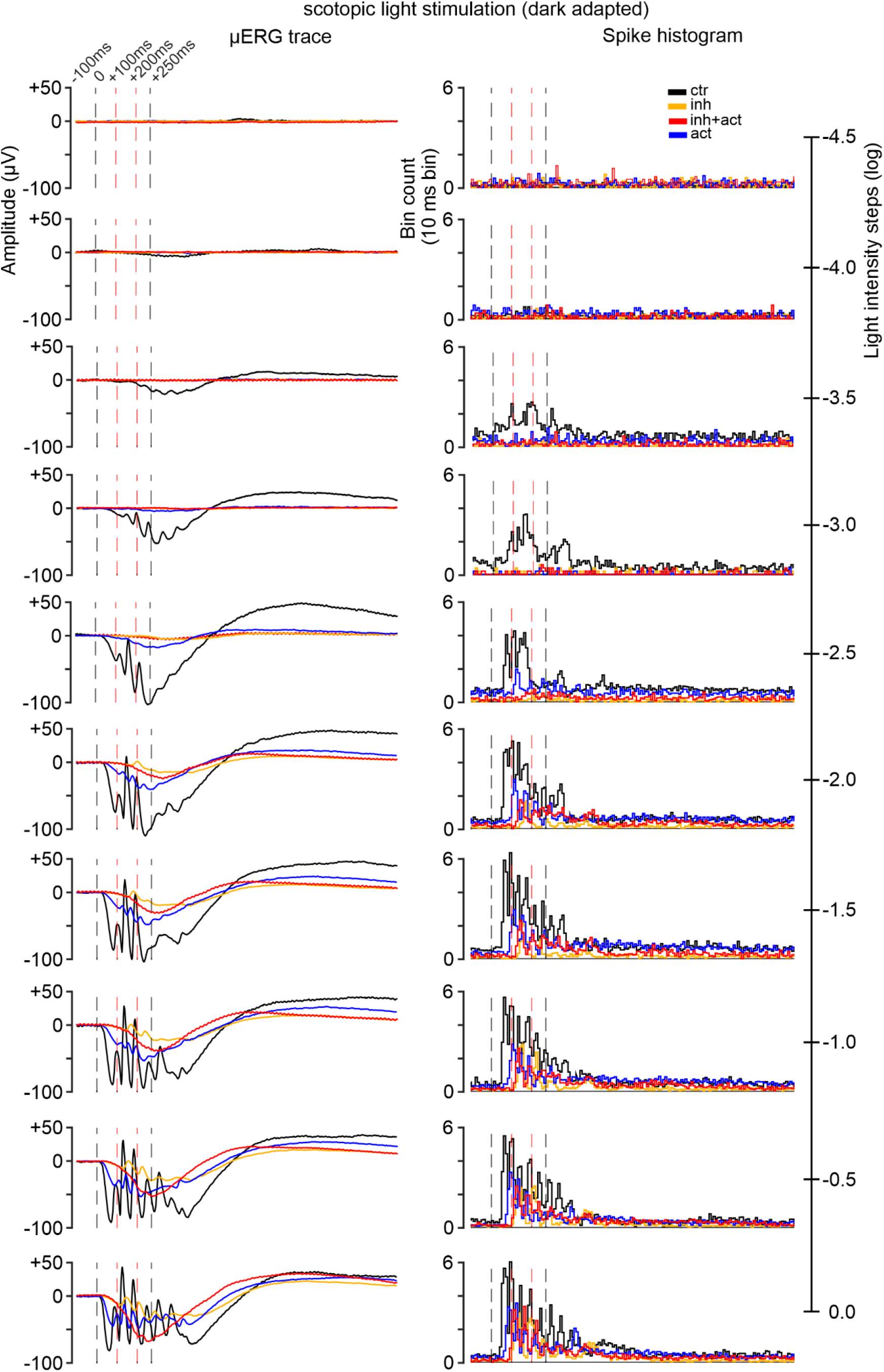
Effects of cGMP analogues on retinal function under scotopic conditions. Representative electrophysiological recordings featuring **A**) the μERG traces and **B**) the corresponding ganglion cells spikes (histogram, 10 ms bin). The summary of this data is presented in Fig. 6. Experimental conditions: control (ctr; black traces), inhibitor (inh; orange), inhibitor and activator combined (inh+act; red) and the activator (act; blue).

**Fig. S4.**
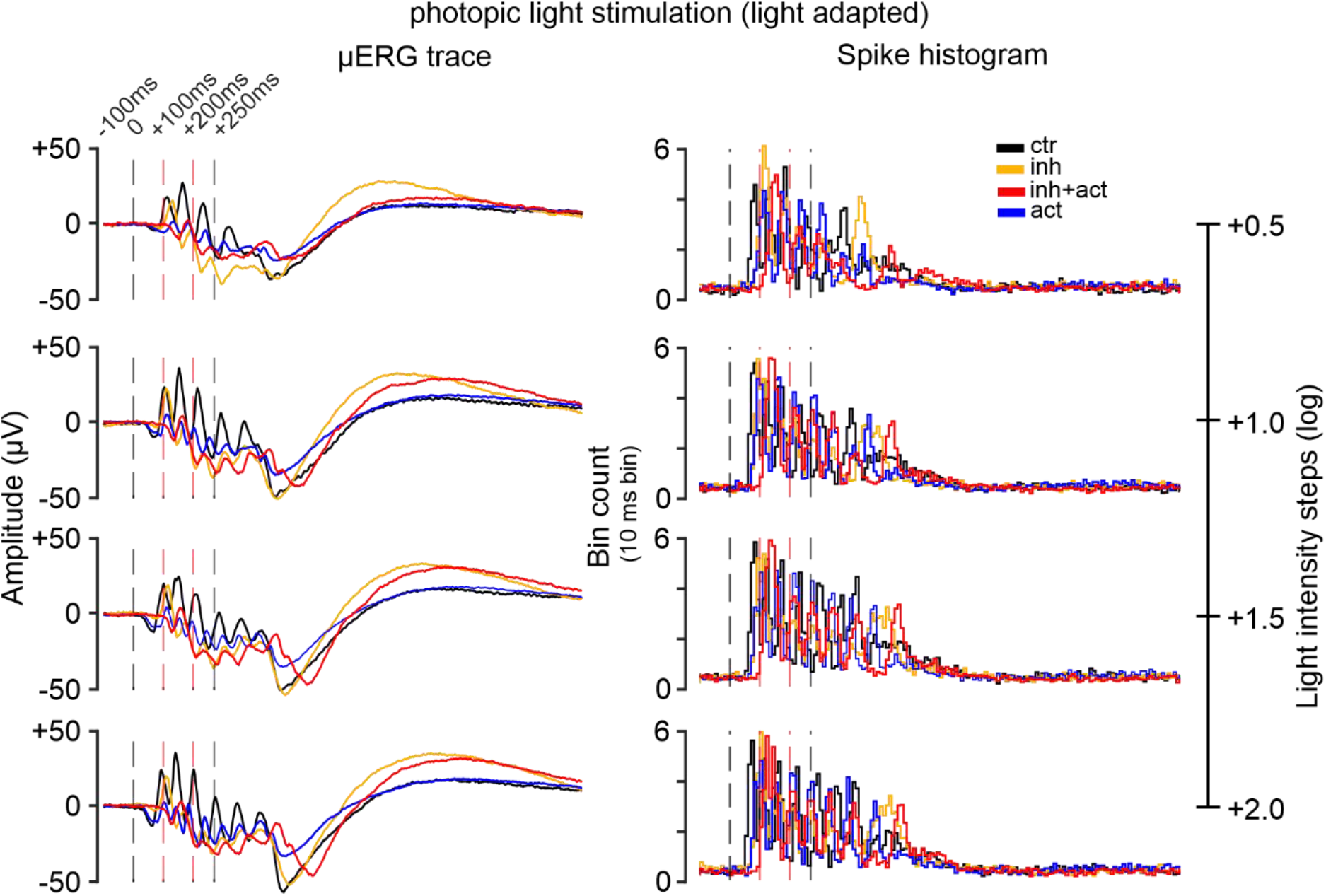
Effects of cGMP analogues on retinal function under photopic conditions. Representative electrophysiological recordings featuring **A**) the μERG traces and **B**) the corresponding ganglion cells spikes (histogram, 10 ms bin). The summary of this data is presented in Fig. 6. Experimental conditions: control (ctr; black traces), inhibitor (inh; orange), inhibitor and activator combined (inh+act; red) and the activator (act; blue).

### Supplementary Tables

**Table S1.**
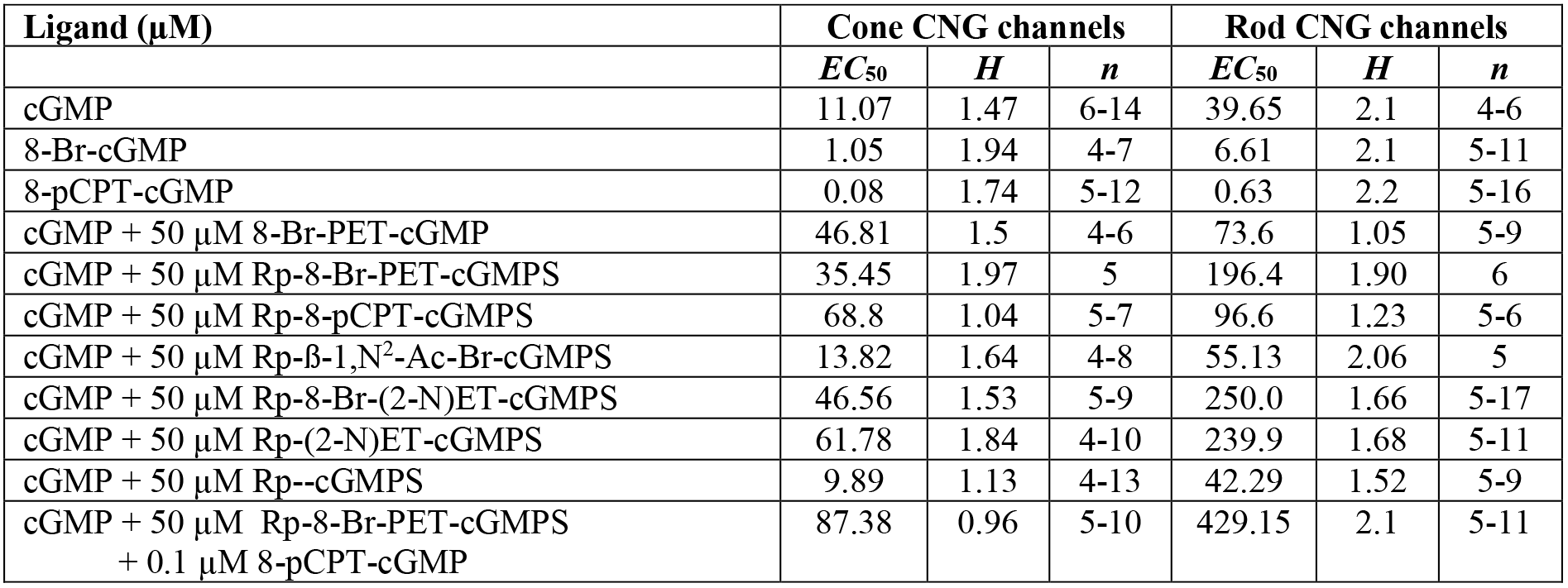
Apparent affinity of retinal CNG channels when activated by either cGMP or combinations of cGMP and cGMP analogues. The table shows the *EC*_50_ values and *H* (Hill coefficient), obtained from the concentrations-activation relationships presented in Figs. 2-4 and Supplementary Fig. 2.

**Table S2.** Effects of the cGMP analogues on the function of the mouse retina under scotopic and photopic stimulation conditions. Presentation of the mean (±std) values for the a-wave amplitude (A) and the corresponding spike responses peaking at a-wave (**B**) from the scotopic and photopic stimulation recordings shown in Fig. 6. Experimental conditions: control (ctr), inhibitor (inh), inhibitor and activator combined (inh+act) and the activator (act).

| A |  | Light intensity steps (log) | Experimental condition |  |  |  |
| --- | --- | --- | --- | --- | --- | --- |
|  |  |  | ctr | inh | inh+act | act |
| a-wave amplitude (μV) | Scotopic | -4.5 | 0.00±0.00 | 0.00±0.00 | 0.00±0.00 | 0.00±0.00 |
|  |  | -4.0 | -0.49±0.78 | 0.00±0.00 | 0.00±0.00 | 0.00±0.00 |
|  |  | -3.5 | -1.00±0.04 | 0.00±0.00 | 0.00±0.00 | 0.00±0.00 |
|  |  | -3.0 | -6.61±2.67 | 0.00±0.00 | 0.00±0.00 | 0.04±0.05 |
|  |  | -2.5 | -29.18±6.75 | 0.00±0.00 | -1.71±0.13 | -3.02±0.74 |
|  |  | -2.0 | -69.70±12.83 | -1.85±0.71 | -3.52±1.96 | -11.59±2.75 |
|  |  | -1.5 | -82.69±12.05 | -2.22±1.40 | -4.36±0.96 | -14.14±3.10 |
|  |  | -1.0 | -87.17±9.94 | -3.62±1.17 | -5.04±2.10 | -18.28±4.38 |
|  |  | -0.5 | -86.51±9.14 | -4.49±1.49 | -7.26±1.94 | -24.30±7.17 |
|  |  | 0.0 | -79.27±8.57 | -7.32±1.37 | -8.55±4.60 | -27.81±6.62 |
|  | Photopic | +0.5 | -5.26±1.66 | -1.53±0.76 | -4.36±3.37 | -3.04±1.68 |
|  |  | +1.0 | -10.36±1.94 | -1.98±0.87 | -9.06±2.42 | -7.72±1.37 |
|  |  | +1.5 | -12.93±2.17 | -1.96±0.89 | -12.55±1.53 | -10.79±1.41 |
|  |  | +2.0 | -14.92±2.15 | -2.27±0.97 | -13.74±3.31 | -12.55±1.72 |
| B |  | Light intensity steps (log) | ctr | inh | inh+act | act |
| Spike count (10 ms bin) | Scotopic | -4.5 | 0.33±0.13 | 1.03±0.27 | 0.92±0.28 | 0.67±0.22 |
|  |  | -4.0 | 0.73±0.00 | 0.98±0.44 | 0.77±0.19 | 0.89±0.07 |
|  |  | -3.5 | 1.79±0.55 | 1.14±0.50 | 0.82±0.15 | 0.86±0.30 |
|  |  | -3.0 | 2.45±0.30 | 0.72±0.10 | 0.71±0.19 | 0.73±0.49 |
|  |  | -2.5 | 3.84±0.44 | 0.83±0.28 | 0.92±0.23 | 1.28±0.51 |
|  |  | -2.0 | 4.57±0.69 | 2.55±0.62 | 1.83±0.33 | 2.41±0.38 |
|  |  | -1.5 | 5.66±0.57 | 2.80±0.63 | 2.13±0.27 | 3.04±0.49 |
|  |  | -1.0 | 5.51±0.54 | 2.91±0.16 | 2.10±0.35 | 3.03±0.52 |
|  |  | -0.5 | 5.49±0.60 | 3.48±0.49 | 2.86±0.25 | 3.60±0.35 |
|  |  | 0.0 | 5.39±0.66 | 3.80±0.26 | 3.04±0.34 | 3.92±0.24 |
|  | Photopic | +0.5 | 4.99±0.45 | 4.85±0.92 | 3.98±1.03 | 4.35±0.79 |
|  |  | +1.0 | 5.36±0.79 | 4.40±0.94 | 4.29±0.85 | 4.43±0.47 |
|  |  | +1.5 | 5.62±0.48 | 4.51±1.07 | 4.50±0.92 | 4.74±1.09 |
|  |  | +2.0 | 5.38±0.87 | 4.92±1.12 | 5.30±1.25 | 4.56±0.92 |

**Table S3.**
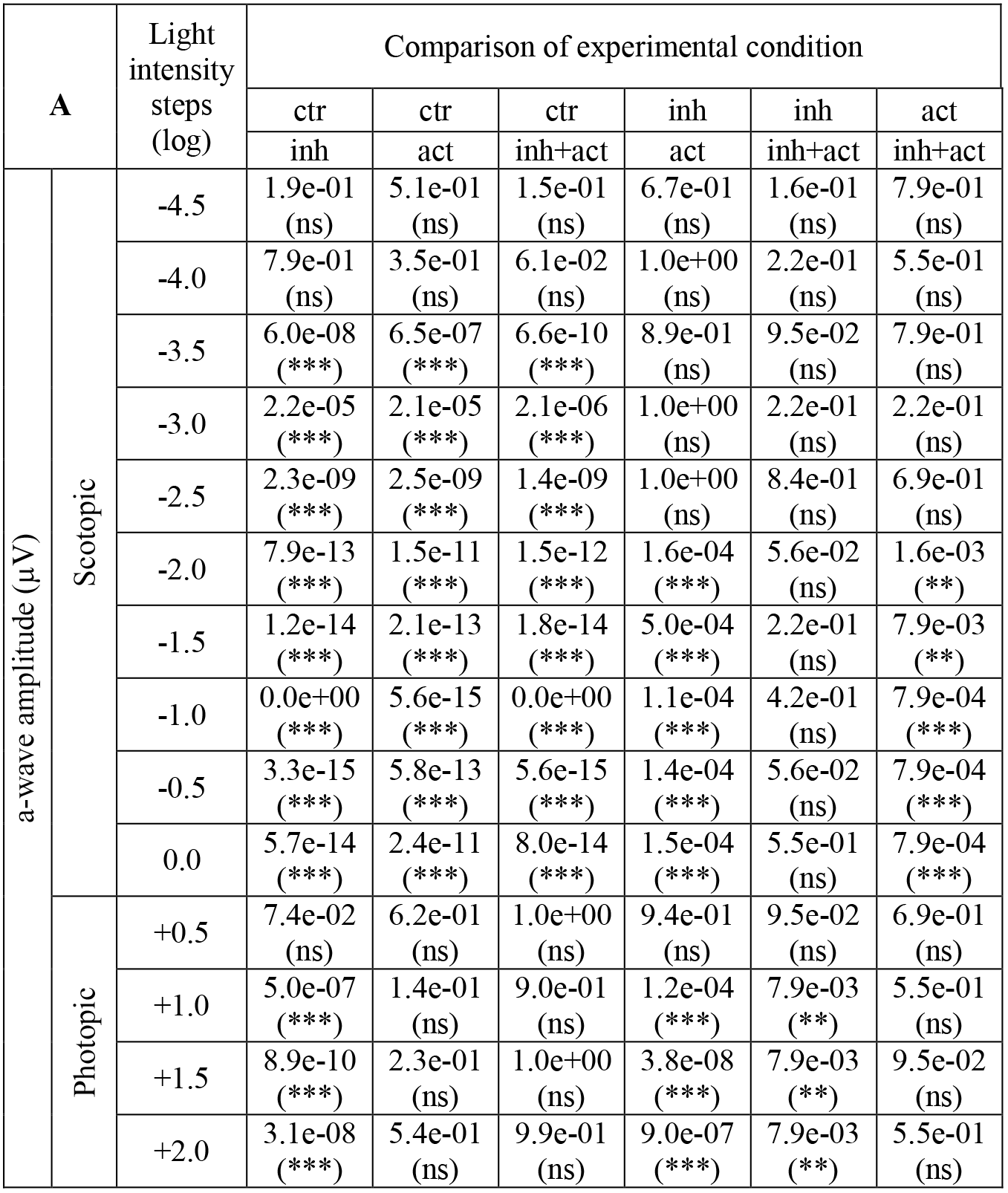
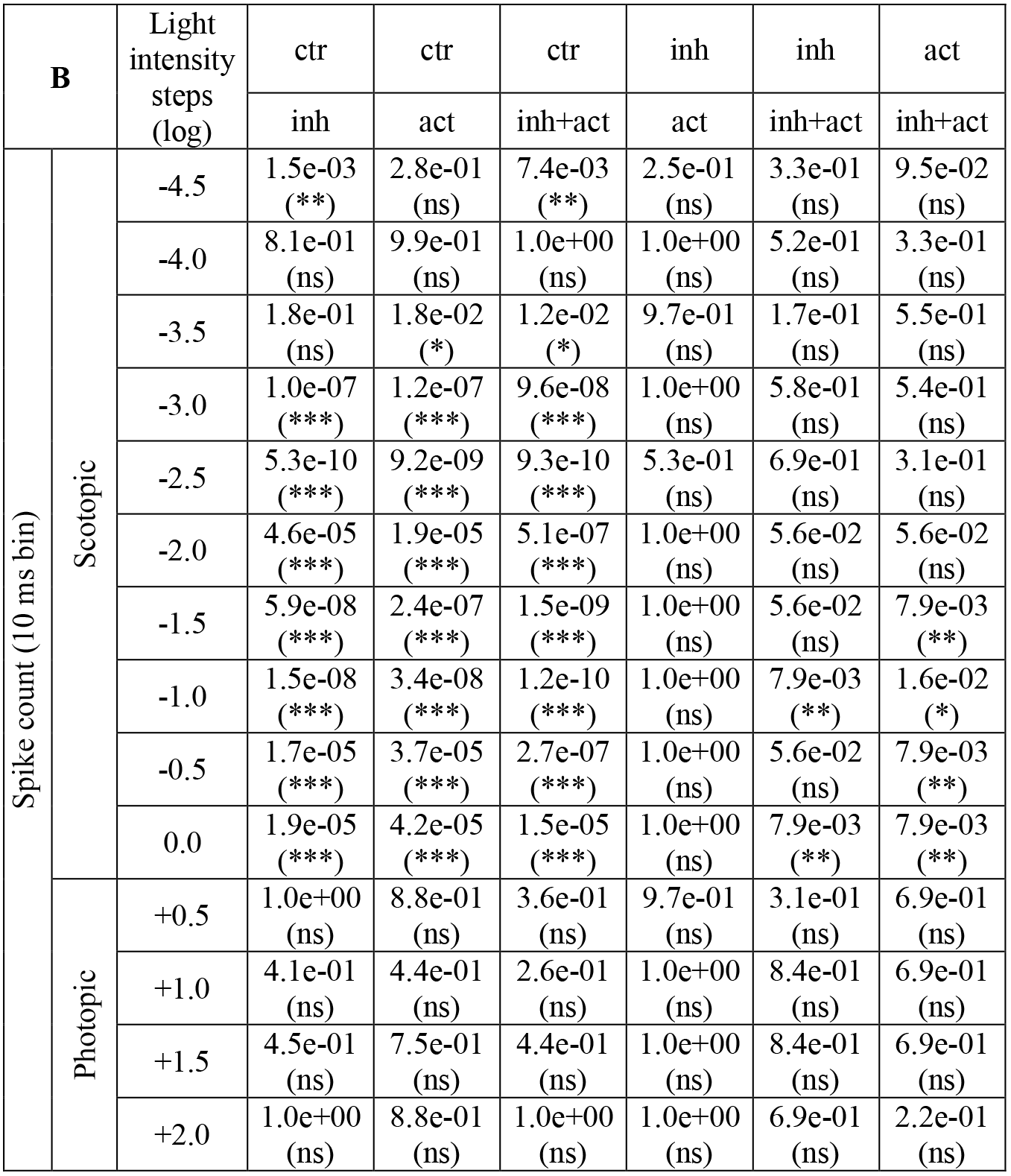
Statistical evaluation of the effects of the cGMP analogues on the function of the mouse retina under scotopic and photopic stimulation conditions. Statistical analysis of the scotopic and photopic stimulation recordings shown in Fig. 6 and Table S2: values of the a-wave amplitude (**A**) and the corresponding spike responses peaking at a-wave (**B**). Statistical significance was estimated by one-way ANOVA followed by the Dunnett’s multiple comparison. Experimental conditions: control (ctr), inhibitor (inh), inhibitor and activator combined (inh+act) and the activator (act).

**Table S4.**
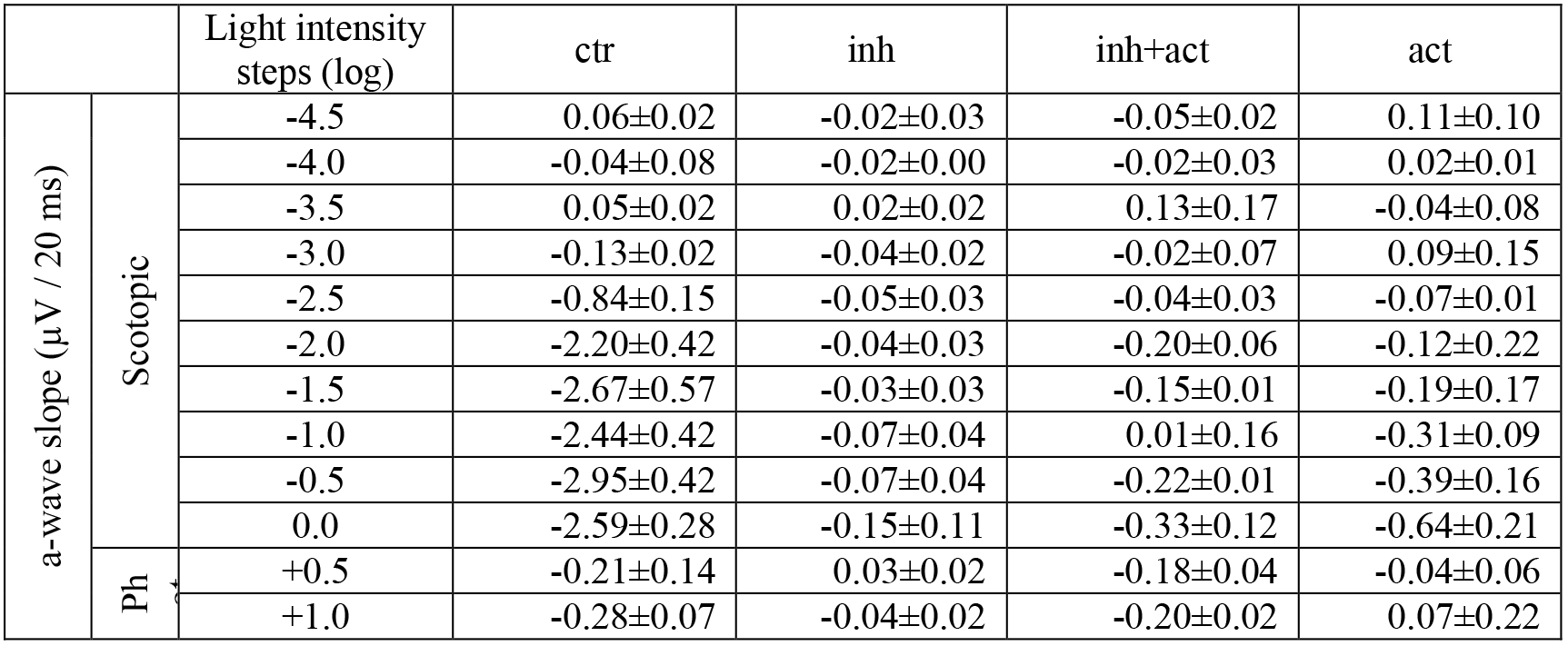
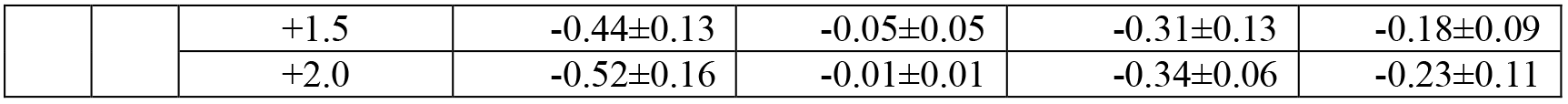
Effects of the cGMP analogues on the temporal response of mouse retina under scotopic and photopic stimulation conditions. Mean (±std) values of the a-wave slope under scotopic and photopic stimulation recordings shown in Fig. 7. Experimental conditions: control (ctr), inhibitor (inh), inhibitor and activator combined (inh+act) and the activator (act).

| a-wave slope (μV / 20 ms) | Light intensity steps (log) | ctr | inh | inh+act | act |
| --- | --- | --- | --- | --- | --- |
|  |  | ctr | inh | inh+act | act |
| Scotopic | -4.5 | 0.06±0.02 | -0.02±0.03 | -0.05±0.02 | 0.11±0.10 |
| | -4.0 | -0.04±0.08 | -0.02±0.00 | -0.02±0.03 | 0.02±0.01 |
| | -3.5 | 0.05±0.02 | 0.02±0.02 | 0.13±0.17 | -0.04±0.08 |
| | -3.0 | -0.13±0.02 | -0.04±0.02 | -0.02±0.07 | 0.09±0.15 |
| | -2.5 | -0.84±0.15 | -0.05±0.03 | -0.04±0.03 | -0.07±0.01 |
| | -2.0 | -2.20±0.42 | -0.04±0.03 | -0.20±0.06 | -0.12±0.22 |
| | -1.5 | -2.67±0.57 | -0.03±0.03 | -0.15±0.01 | -0.19±0.17 |
| | -1.0 | -2.44±0.42 | -0.07±0.04 | 0.01±0.16 | -0.31±0.09 |
| | -0.5 | -2.95±0.42 | -0.07±0.04 | -0.22±0.01 | -0.39±0.16 |
| | 0.0 | -2.59±0.28 | -0.15±0.11 | -0.33±0.12 | -0.64±0.21 |
| Ph | +0.5 | -0.21±0.14 | 0.03±0.02 | -0.18±0.04 | -0.04±0.06 |
| | +1.0 | -0.28±0.07 | -0.04±0.02 | -0.20±0.02 | 0.07±0.22 |

|  |  |  |  |  |  |  |
| --- | --- | --- | --- | --- | --- | --- |
|  |  | +1.5 | -0.44±0.13 | -0.05±0.05 | -0.31±0.13 | -0.18±0.09 |
|  |  | +2.0 | -0.52±0.16 | -0.01±0.01 | -0.34±0.06 | -0.23±0.11 |

**Table S5.** Statistical evaluation of the effects of the cGMP analogues on the response kinetics of mouse retina under scotopic and photopic stimulation conditions. Statistical significance of the a-wave slope under scotopic and photopic stimulation recordings shown in Fig. 7 and Table S4. The slope was calculated only for a-wave amplitudes exceeding the control base line indicating a retinal light response. Statistical significance was estimated by one-way ANOVA followed by the Dunnett’s multiple comparison. Experimental conditions: control (ctr), inhibitor (inh), inhibitor and activator combined (inh+act) and the activator (act).

|  |  | Light intensity steps (log) | ctr | ctr | ctr | inh | inh | act |
| --- | --- | --- | --- | --- | --- | --- | --- | --- |
|  |  |  | inh | act | inh+act | act | inh+act | inh+act |
| a-wave slope (μV / 20 ms) | Scotopic | -4.5 | NaN | NaN | NaN | NaN | NaN | NaN |
|  |  | -4.0 | NaN | NaN | NaN | NaN | NaN | NaN |
|  |  | -3.5 | NaN | NaN | NaN | NaN | NaN | NaN |
|  |  | -3.0 | NaN | NaN | NaN | NaN | NaN | NaN |
|  |  | -2.5 | NaN | 2.0e-12 (***) | 9.4e-13 (***) | NaN | NaN | 9.5e-02 (ns) |
|  |  | -2.0 | 8.9e-12 (***) | 1.7e-11 (***) | 3.6e-11 (***) | 1.0e+00 (ns) | 7.9e-03 (**) | 8.4e-01 (ns) |
|  |  | -1.5 | 1.0e-11 (***) | 3.1e-11 (***) | 2.4e-11 (***) | 9.9e-01 (ns) | 7.9e-03 (**) | 6.9e-01 (ns) |
|  |  | -1.0 | 6.4e-13 (***) | 4.7e-12 (***) | 3.4e-13 (***) | 5.6e-01 (ns) | 5.5e-01 (ns) | 7.9e-03 (**) |
|  |  | -0.5 | 8.9e-15 (***) | 8.2e-14 (***) | 2.4e-14 (***) | 1.9e-01 (ns) | 7.9e-03 (**) | 1.5e-02 (*) |
|  |  | 0.0 | 1.2e-14 (***) | 8.6e-13 (***) | 5.2e-14 (***) | 2.1e-03 (**) | 5.6e-02 (ns) | 3.2e-02 (*) |
|  | Photopic | +0.5 | 1.8e-03 (**) | 4.5e-02 (*) | 1.0e+00 (ns) | 8.6e-01 (ns) | 7.9e-03 (**) | 7.9e-03 (**) |
|  |  | +1.0 | 2.6e-02 (*) | 5.2e-04 (***) | 9.5e-01 (ns) | 6.8e-01 (ns) | 7.9e-03 (**) | 7.9e-03 (**) |
|  |  | +1.5 | 1.5e-05 (***) | 3.0e-03 (**) | 3.1e-01 (ns) | 2.4e-01 (ns) | 7.9e-03 (**) | 1.5e-01 (ns) |
|  |  | +2.0 | 3.8e-07 (***) | 8.8e-04 (***) | 6.7e-02 (ns) | 1.4e-02 (*) | 7.9e-03 (**) | 2.2e-01 (ns) |

## Acknowledgments

We thank S. Bernhardt, K. Schoknecht, A. Kolchmeier, Uta Enke and Claudia Ranke from Institute of Physiology II (Jena) for excellent technical assistance. We thank also J. Kusch and K. Benndorf from Institute of Physiology II (Jena) for excellent comments on the manuscript.

We thank U. Manzau from Biolog LSI GmbH & Co. KG for expert technical assistance during synthesis and purification of cGMP analogues.

We thank also Prof. Eberhardt Zrenner from the Institute for Ophthalmic Research, University of Tübingen, for excellent comments on the manuscript.

## Author contributions

SW and SK studied the effect of the cGMP analogues on heterologously-expressed CNG channels; VP studied the effect of the cGMP analogues on the gating kinetics of CNG channels; CM performed the molecular biology work; AR and FS were responsible for the design of cGMP analogues. AR performed synthesis of cGMP analogues. FS supervised synthesis and purification of cGMP analogues; VN designed the CNG-channel experiments, performed electrophysiological measurements, analyzed/ interpreted the electrophysiological data; WH, SD, FPD designed the μERG experiments; WH performed electrophysiological measurements, analyzed/ interpreted the electrophysiological data. VN, WH and FPD prepared the figures and wrote the manuscript in collaboration with all coauthors.

## Data and materials availability

All data are available in the main text or the supplementary materials.

